# *In silico* learning of tumor evolution through mutational time series

**DOI:** 10.1101/577171

**Authors:** Noam Auslander, Yuri I. Wolf, Eugene V. Koonin

## Abstract

Cancer arises through the accumulation of somatic mutations over time. Understanding the sequence of mutation occurrence during cancer progression can assist early and accurate diagnosis and improve clinical decision-making. Here we employ Long Short-Term Memory networks (LSTMs), a class of recurrent neural network, to learn the evolution of a tumor through an ordered sequence of mutations. We demonstrate the capacity of LSTMs to learn complex dynamics of the mutational time series governing tumor progression, allowing accurate prediction of the mutational burden and the occurrence of mutations in the sequence. Using the probabilities learned by the LSTM, we simulate mutational data and show that the simulation results are statistically indistinguishable from the empirical data. We identify passenger mutations that are significantly associated with established cancer drivers in the sequence and demonstrate that the genes carrying these mutations are substantially enriched in interactions with the corresponding driver genes. Breaking the network into modules consisting of driver genes and their interactors, we show that these interactions are associated with poor patient prognosis, thus likely conferring growth advantage for tumor progression. Thus, application of LSTM provides for prediction of numerous additional conditional drivers and to reveal hitherto unknown aspects of cancer evolution.

**Significance:** Cancer is caused by the effects of somatic mutations known as drivers. Although a number of major cancer drivers have been identified, it is suspected that many more comparatively rare and conditional drivers exist, and the interactions between different cancer-associated mutations that might be relevant for tumor progression are not well understood. We applied an advanced neural network approach to learn the sequence of mutations and the mutational burden in colon and lung cancers, and to identify mutations that are associated with individual drivers. A significant ordering of driver mutations is demonstrated, and numerous, previously undetected conditional drivers are identified. These findings broaden the existing understanding of the mechanisms of tumor progression and have implications for therapeutic strategies.

## Introduction

Tumorigenesis is a multistep process characterized by accumulation of somatic mutations, which contribute to tumor growth, clinical progression, immune escape and the development of drug resistance (1,2). The somatic mutations found in a tumor cell are accumulated over the lifetime of the cancer patient, so that some mutations are acquired in early steps of tumorigenesis and even in pre-malignant cells (3). Relatively small subsets of these mutations are established tumor drivers, whereas the remainder are thought to be passengers that do not confer growth advantage or may even negatively affect tumor fitness (4–7). Although molecular and cell biology studies have revealed many mechanistic details of tumorigenesis (4,8–10), our understanding of the tumor evolution dynamics remains limited, presumably, due to the complexity of the process and the abundance of passenger events that could be randomly distributed but might exert various effects on tumor fitness and properties (11,12).

In colorectal cancer, tumor development has been explained by a multistep model of carcinogenesis that describes the progression of a benign adenoma to a malignant carcinoma through a series of well-defined histological stages that are linked to a mutational time series, i.e. the temporal sequence of occurrence of driver mutations (13–16). Similar stepwise models have been developed for other types of adenocarcinomas (17,18) although the temporal succession of molecular changes characterizing the progression of these tumors has not been elucidated at the level of confidence it has for colon cancer (19). Given that the somatic alterations in some tumors can be represented as a multistep sequence of events, we conjectured that time series learning algorithms could be applied to the sequence of somatic mutations, potentially revealing the complex dynamics of the tumor evolution and enabling a variety of context-specific predictions.

To this end, we employed Long Short Term Memory (LSTM) networks (20), a type of Recurrent Neural Networks (21) capable of learning long-term dependencies in a time series sequence. These networks have achieved major success in time-series prediction tasks and for learning evolution of recurrent systems (20,22–24). We demonstrate here the utility of applying LSTM to time-ordered mutational data in colon and lung adenocarcinomas. We define a pseudo-temporal gene ranking, representing each tumor sample as a binary text, which can be used for training LSTM in a similar manner to that applied in naturel language texts classification (25–27). Applied to a discrete version of the mutational data, ordered into an approximate temporal sequence, these networks can be used to predict the mutational load from a limited number of mutations, and for stepwise prediction of the occurrence of following mutations in the series. Using the sequence dynamics learned by the models, we reconstruct sequences of mutations that are statistically indistinguishable from the original observations. We find that the occurrence of distinct subsets of mutations could be predicted from the timeline of the major cancer drivers. Investigation of the driver genes contributing to the prediction of each of these mutations uncovers numerous driver-interactor gene pairs that are highly and specifically enriched with different types of independently identified interactions. We further derive modules of driver genes and find that their interactions are associated with poor survival rate, suggesting that the driver and associated passengers jointly promote tumor growth.

## Results

Throughout this work, we use a discretized version of the mutational data from two tumor types, namely, colon cancer, where the stepwise evolutionary model has been established (13–16), and lung cancer, where such a model has been suggested but not widely accepted as it is in the case of colon cancer (17,19). The “snapshot” mutational datasets are ordered into an approximate temporal sequence that is estimated via the training sets for each tumor type. For each classification task, we trained LSTM networks using The Cancer Genome Atlas (TCGA) (28) time-ordered mutational data for these tumor types, and tested the network performance using independent data sets (Table 1).

**Table 1.** Datasets used for training and testing, for colon and lung adenocarcinomas.

|  | Colon | Lung |
| --- | --- | --- |
| Training | COAD (n = 253) | LUAD (n=506) |
| Test | Colorectal Adenocarcinoma (DFCI, Cell Reports 2016 (67), n = 612)<br>Colorectal Adenocarcinoma (Genentech, Nature 2012 (68), n=72) | Lung Adenocarcinoma (Broad, Cell 2012 (69), n = 183) |

### Predicting tumor mutational load from the mutational time series

We aim to represent tumor mutations as a discrete timeline of mutational events such that the appearance of a mutation in the timeline would correspond to their estimated place in the tumor evolution. We thus calculate a score for every mutation as the ratio of its occurrences in the presence and in the absence of other mutations (see Methods for details). This form of the score is motivated by the assumption that mutations that appear late in the tumor evolution are more likely to be fixed when other mutations are present, supported by a previously established notion that co-occurring events more often take place at late stages in tumor evolution (29–32). Sorting the mutations by the scores evaluated on our training datasets, we find that this estimated order of mutation appearance is in agreement with the established succession of the key drivers in colon adenocarcinoma (SI Appendix Fig. S1A). In particular, the APC mutation is an early driver event, followed by CTNBB1, KRAS and SMAD4, whereas PIC3CA and TP53 mutations occur later in tumor development (13,33). Moreover, we find that the scores significantly correlate with the ratio between the frequency of a mutation in colon adenomas and its frequency in colon carcinomas (from COSMIC database (34,35), SI Appendix, Fig. S1B), further supporting the relevance of this score for the inference of the order of mutations.

We then used the time-ordered mutational data to predict the overall mutational load in the respective tumors and evaluate the number of mutations required for an accurate prediction. To this end, we trained LSTM networks aiming to predict ‘high’ vs. ‘low’ mutational load (Methods) from a time series of mutations. Given a discrete time series of mutation occurrences, the LSTM network is trained using the series up to a time *t*, and is applied to the left-out test to produce scores that reflect the probability of each sample in the test to have a ‘high’ mutational load. Starting from the final time point (the last mutation in the series, likely occurring late in the tumor evolution), we find that prediction of the mutational load saturates with high performance (AUC>0.95) with less than 100 mutations for both colon and lung adenocarcinomas (Fig. 1A). The sets of genes that contribute to the mutational load prediction significantly overlap between colon and lung cancers (P-value ∼= 0 for the last 100 genes). Notably, this overlap includes genes that encode some of the longest human proteins that perform various organizing roles in either intracellular or inter-cellular interactions, such as titin (TTN), mucin-16 (MUC16) and nesprin-1 (SYNE1). TTN is one of the most commonly mutated early drivers in colon cancer (36). MUC16 is implicated in the progression of several cancers, apparently, via interactions with the immune system, and is emerging as an important target for cancer therapy (37). SYNE1, a cytoskeleton organizer, although less thoroughly characterized, appears to contribute to DNA damage response, and thus, to genome instability and tumorigenesis (38). Thus, the genes that contribute to mutation load prediction in both types of cancer seem to reveal common biological themes. Moreover, the scores assigned by the LSTM using only the 20 latest mutations (i.e. the 20 mutations with the highest order scores) as a sequence are highly correlated with the observed mutational load in all test sets (Fig. 1B-D). Using the time series from the earliest time point, however, results in almost random prediction performance with similar number of mutations (SI Appendix, Fig. S2), suggesting that the ultimate mutational burden of a tumor depends primarily on mutations that occur late in tumor evolution. Training linear classifiers to predict the mutational load from the same sequences of mutations, or randomly selected mutations (see Methods) resulted in significantly inferior performance compared to the LSTM (Fig. 1E-G), suggesting that the LSTM networks learn complex dynamics within the mutational data that could not be captured by conventional classification approaches.

**Fig. 1.**
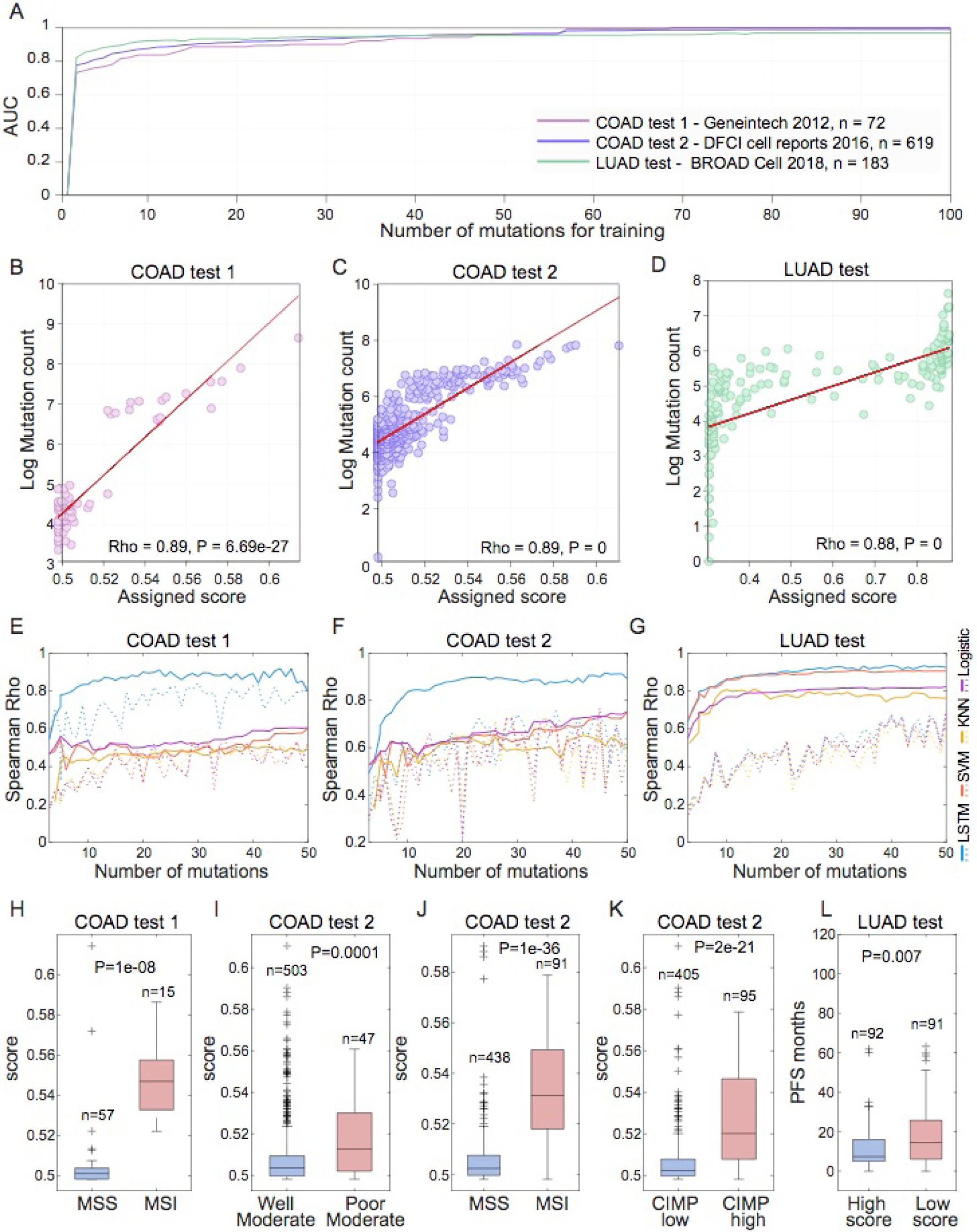
Prediction of tumor mutational load from the mutational time series. (A) The test AUCs (Y-axis) obtained for training LSTMs on different lengths of mutation sequences (X-axis) when starting from the latest ordered mutation, for colon and lung test sets. (B)-(D) Correlation between the score assigned by LSTMs with the 20 latest mutations in the time series (x-axis), and the true mutational load (y-axis, log transformed). (E)-(G) Spearman correlation between the scores assigned by different learning models and the observed mutational load (y-axis) when using different number of mutations from these that are ordered latest in the sequence (up to 50 mutations, x-axis). The dashed lines show the results for classifiers trained on randomly selected mutations (rather than the ordered sequence of mutations that is shown by solid lines). (H)-(K) Scores assigned by LSTM using the last 20 mutations for colon cancer patients with different clinical characteristics. (L) Progression-free survival (PFS) of lung cancer patients in the test set, of samples assigned with high vs. low (using the median) scores with the last 20 mutations. Abbreviations: COAD, Colon adenocarcinoma, LUAD, Lung adenocarcinoma

The scores assigned to predict the mutational load are also associated with clinical phenotypes. For the colon cancer test sets, the scores assigned by LSTM trained with the 20 latest mutations in the sequence are significantly higher in samples of primary tumors assigned with poor grade vs. moderate grade, in high vs. low microsatellite instability and in high vs. low CpG island methylation phenotypes (Fig. 1H-K). In the lung cancer test set, higher scores are associated with lower progression-free survival (PFS, Fig. 1L).

### Predicting occurrence of mutations in the sequence and simulated data analysis

We then explored the possibility of predicting the occurrence of mutations in the course of tumor evolution from other mutations in the time series. To this end, the LSTM networks were trained to predict the occurrence of each mutation *M*_*t*_ in the time series, using the time series of mutations from the latest mutation up to the *M*_*t+1*_ mutation (thus, predicting earlier mutations from later ones). The occurrence of most mutations in the sequence could be predicted with good accuracy, with the median AUC = 0.88, 0.73 and 0.69 for the first and second colon test sets, and for the lung test set, respectively (Fig. 2A). Mutations for which occurrence could not be predicted (AUC < 0.5, 2% and 17% of the mutations in colon and lung cancers, respectively) were significantly less common than those that were readily predictable (Rank-sum P-value = 0.001 for colon and 2.4e-106 for lung, SI Appendix, Table S1), implying that the occurrence of these low frequency mutations is not linked to tumor progression.

**Fig. 2.**
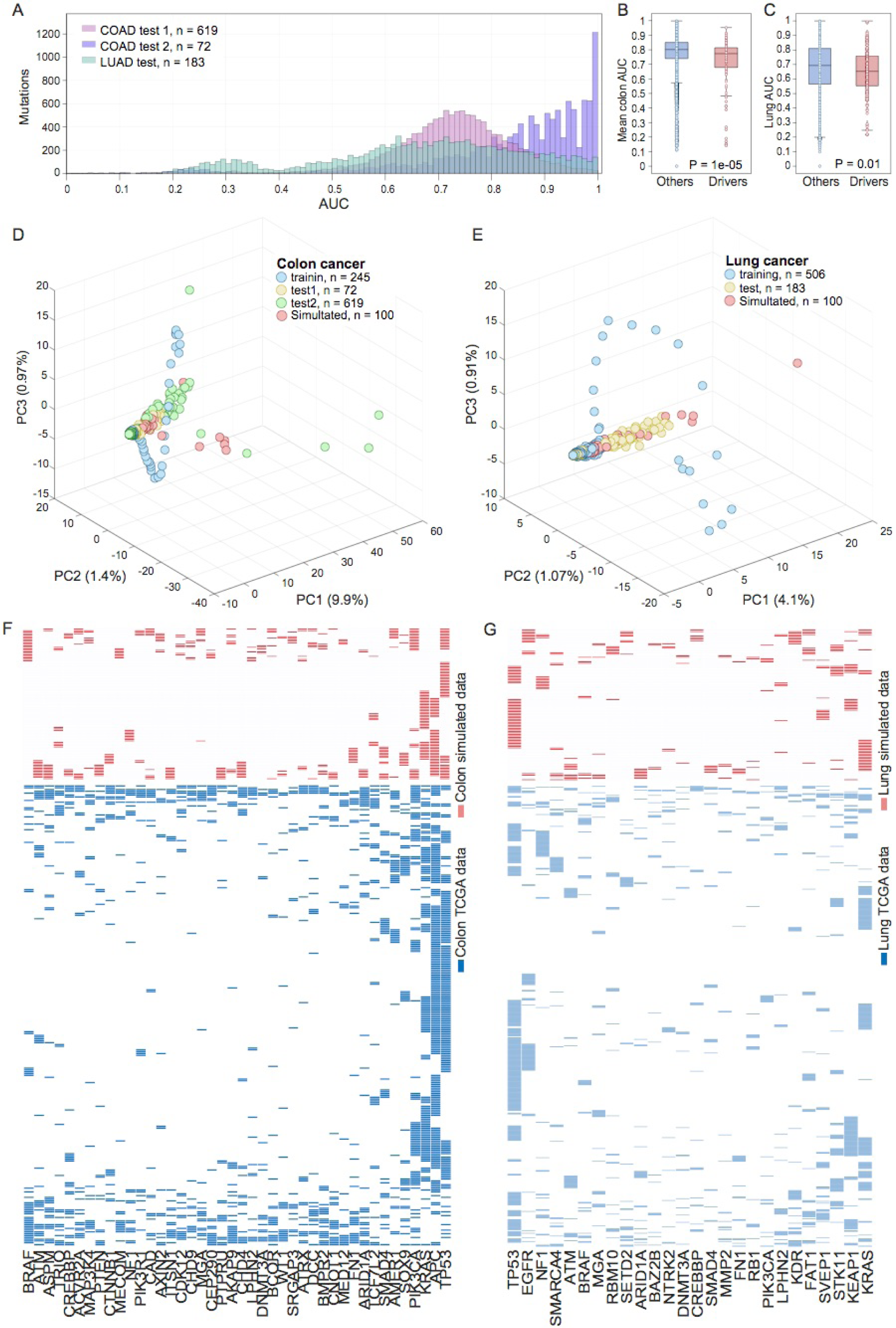
Prediction and generation of the sequence of mutations. (A) Histogram of mutations count (Y-axis) for each performance level (AUC, X-axis) for mutation prediction in a sequence for colon cancer test sets (pink and purple bars for test sets 1 and 2, respectively), and for lung cancer test set (green bars). (B) Mean AUC of mutation prediction in the sequence for the two colon test sets: comparison of drivers with all other genes. (C) AUC of mutation prediction in the sequence for the lung test set: comparison of drivers with all other genes. (D)-(E) Scatter plots of PC1-PC3 obtained by PCA applied to the combined mutational data from all datasets used and the simulated samples, for colon and lung cancers, respectively. The percentage of variance explained by each PC is indicated in parentheses. (H)-(K) Presence-absence patterns for the high frequency cancer drivers in the reconstructed mutational samples (red) and the TCGA mutational data (blue) for colon and lung cancers, respectively. The samples are ordered by the hierarchical clustering results, with the Euclidean distance metric and average linkage.

More unexpected than the poor prediction for low frequency mutations, the prediction accuracy for mutations in known cancer driver genes was significantly lower than that for mutations in all other genes, in both tumor types (Fig. 2B-C). This observation is maintained when including only genes that are frequently mutated (SI Appendix, Fig. S3). These findings indicate that the driver mutations that determine the course of tumorigenesis could not be readily predicted from the rest of the mutational landscape of a given tumor type. In contrast, many passenger mutations tend to be linked to specific drivers (39) and thus could be predicted with confidence. Notably, however, the subset of driver mutations for which good prediction accuracy was achieved, included DNA repair genes, such as MLH1, MSH2, MSH6, PMS2 and MUTYH for colon cancer, and DDX5 and CHD1L for lung cancer, conceivably, due to their effect on other mutations in the sequence (SI Appendix, Table S1).

Recently, it has been shown that LSTM recurrent neural networks can be used to generate complex sequences by simply predicting one data point at a time (24). We hence reasoned that similar technique could be utilized to generate the mutational time sequence, and thus could be employed for mutational data reconstruction. We used the LSTM scores to predict the occurrence of each mutation in the time series (from the latest to the earliest), to determine the occurrence of one mutation at a step in a simulated time series (see Methods for details). In this manner, we reconstructed 100 mutational samples for colon cancer, and 100 samples for lung cancer (SI Appendix, Tables S2-3). The K-means clustering analysis did not separate the simulated data from the real data (SI Appendix, Fig. S4A-B), and Principal Component Analysis (PCA) showed notable similarity between the simulated datasets and the real ones (Fig. 2D-E), which is maintained when considering only frequently mutated genes (SI Appendix, Fig. S5). This similarity was observed also when using the tSNE dimensionality reduction method (40) (SI Appendix, Fig. S4C-D). Considering the patterns of occurrence of the frequently mutated cancer drivers (those with the frequency of mutation in the top 10%) in the simulated data, we identified clear similarities to the observed patterns in the original data, such as the mutual exclusion of APC, KRAS and TP53 in colon cancer, and of KRAS and MGA in lung cancer (Fig. 2F-G, Fig. S6).

To evaluate the effect of the actual order of the mutations in the sequence function on the prediction of the preceding mutations, we repeated this analysis for the 300 last mutations, in both colon and lung cancers, after randomly permuting the order of mutations in the sequence up to each predicted mutation (10 random permutations for each prediction task). The comparison of the results with those obtained with the original, ordered sequence of mutations shows a dramatic drop in the prediction accuracy (paired rank-sum P-value<0.06, 0.02 and 9e-6 for colon test set1, colon test set 2 and lung test set, respectively, SI Appendix, Fig. S7). This is a strong indication that the actual order of mutations is important for predicting the mutational sequence.

### Predicting associations between major cancer drivers and other genes

Given that most mutations are predicted with high accuracy from the time sequence, we investigated the possibility that some mutations (currently classified as passenger) might also be predicted from the ordered time sequence of mutations in cancer driver genes which play key roles in tumor development. Mutations that might be predicted through their association with major cancer drivers are of obvious interest because they could potentially contribute to different aspects of tumorigenesis. To explore such potential associations, we selected well characterized, major cancer drivers in each tumor type studied (n=42 for colon and n=26 for lung; see Methods) and utilized the discrete time series of their occurrences to predict the occurrence of other mutations. This analysis yielded 354 genes for colon cancer and 273 genes for lung cancer in which mutations could be predicted robustly with high AUC from multiple points in the time series of the major drivers (Methods, SI Appendix, Table S4).

Given that these mutations are accurately predicted using a short mutational sequence that includes only the major drivers, we hypothesized that the respective genes interact with these drivers. To further characterize the potential functional connections between the major drivers and the identified associated genes, we first determined which driver contributed to the prediction of each of the identified driver-associated mutations (see Methods) to generate a list of driver-interactor pairs for colon and lung cancers (Fig. 3A, SI Appendix, Table S5). A STRING interactions enrichment analysis (41,42) (see Methods) showed that, for 68 and 39 predicted driver-interactor pairs in colon and lung cancers, respectively, the interaction is validated under the multiple criteria implemented in STRING (hyper-geometric P-value∼=0 and 5.2675e-04 for colon and lung, respectively). When the major cancer drivers were analyzed individually, we found significant (P-value <0.05) STRING enrichment between about 22% of both colon and lung cancer major drivers and their predicted interactors, a result that is unlikely to be obtained by chance (Fig. 3A, permutation P-value <0.001 for both colon and lung).

**Fig. 3.**
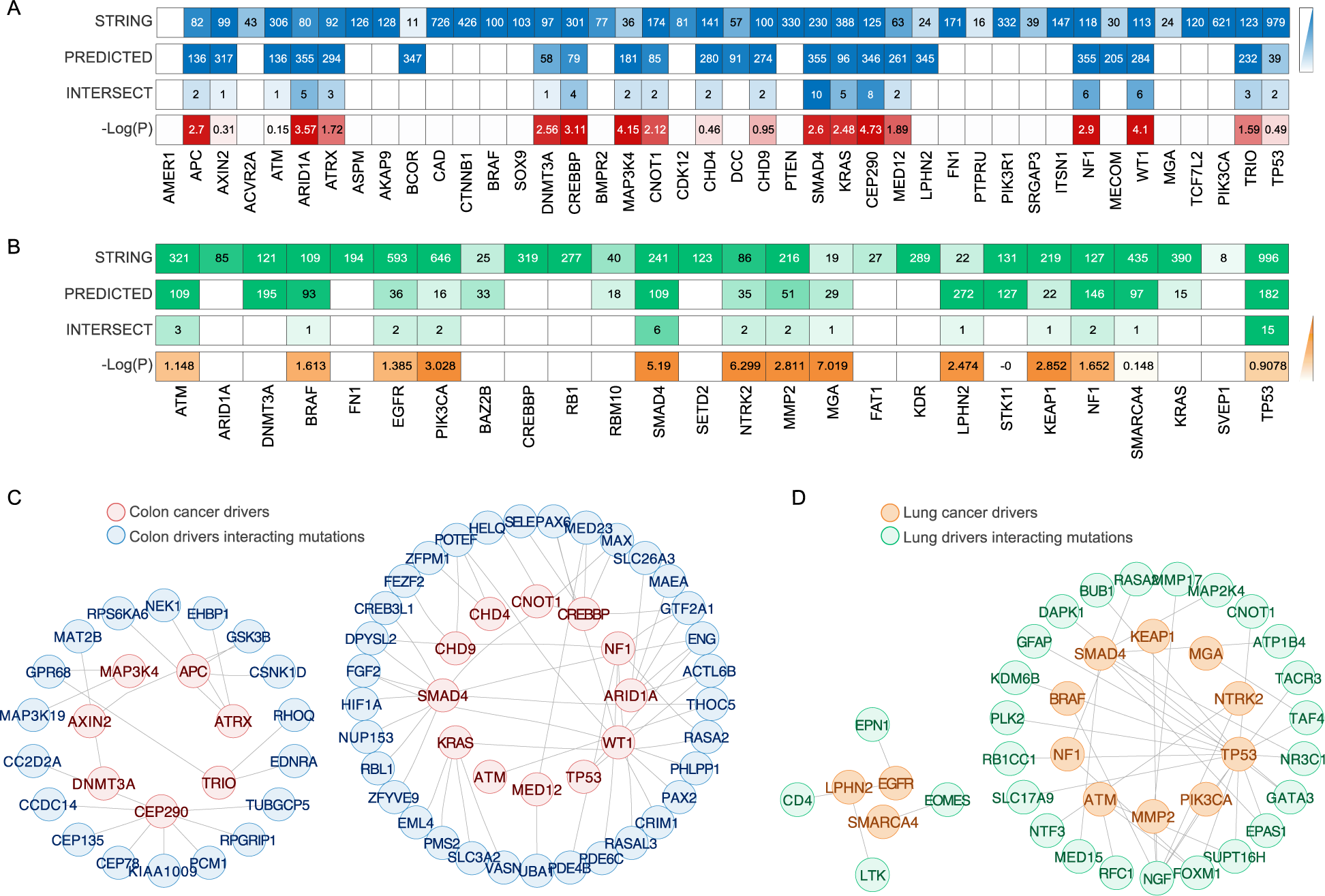
STRING-validated interactions of major drivers in colon and lung cancers. (A)-(B) Heatmaps showing, for each major colon and lung cancer driver, respectively, the number of STRING interactions within the mutational data (top panel), the number of predicted interactions (second to top panel), the number of interactions in the intersection (third to top panel) and the hyper-geometric P-value (bottom panel). (C)-(D) The networks of STRING-validated interactions for colon and lung cancers, respectively.

The node degrees of the major cancer drivers in the network of STRING-validated interactions vary substantially within the networks inferred for each tumor type and differ between the colon and lung networks for shared drivers, with the mean degree of 3.5 for colon and 3 for lung (Fig. 3C-D). In both networks, SMAD4, a gene encoding a protein involved in the TGF-beta signaling pathway (43), is highly connected. Most of the genes connected with SMAD4 in these networks are also involved in TGF-beta signaling but the specific subsets of these genes differ between the colon and lung networks. Several of these interactions have been reported previously. In particular, MAP2K4 has been identified as a conditional tumor suppressor in lung adenocarcinomas but not in colon cancer (44), whereas mutations in HIF1A are associated with the worst prognosis in colon cancer (45); furthermore, HIF1A protein physically interacts with SMAD4 under hypoxic conditions in colon cancer cell lines (46). The degree of TP53 is considerably higher in the lung cancer network than in the colon cancer network, possibly, due to different TP53 mutants that are observed in these tumors (47) that have different interacting partners (48). Notably, in both the colon and the lung networks, TP53, SMAD4, ATM and NF1 belong to the same strongly connected network module, whereas major tumor-specific cancer drivers such as APC (colon) and EGFR (lung) are disconnected.

Next, we systematically assessed whether the predicted interactors of the major cancer drivers are involved in the same or similar processes with the corresponding drivers. Using GO enrichment (49,50), we find that, for both colon and lung cancers, all major drivers with predicted interactors share a significant (with hyper-geometric P-value<0.05) overlap of the sets of GO processes with their predicted interactors (Fig. 4A-B). This level of enrichment in shared processes is unlikely to be reached by chance (Permutation P<0.001 for both colon and lung).

**Fig. 4.**
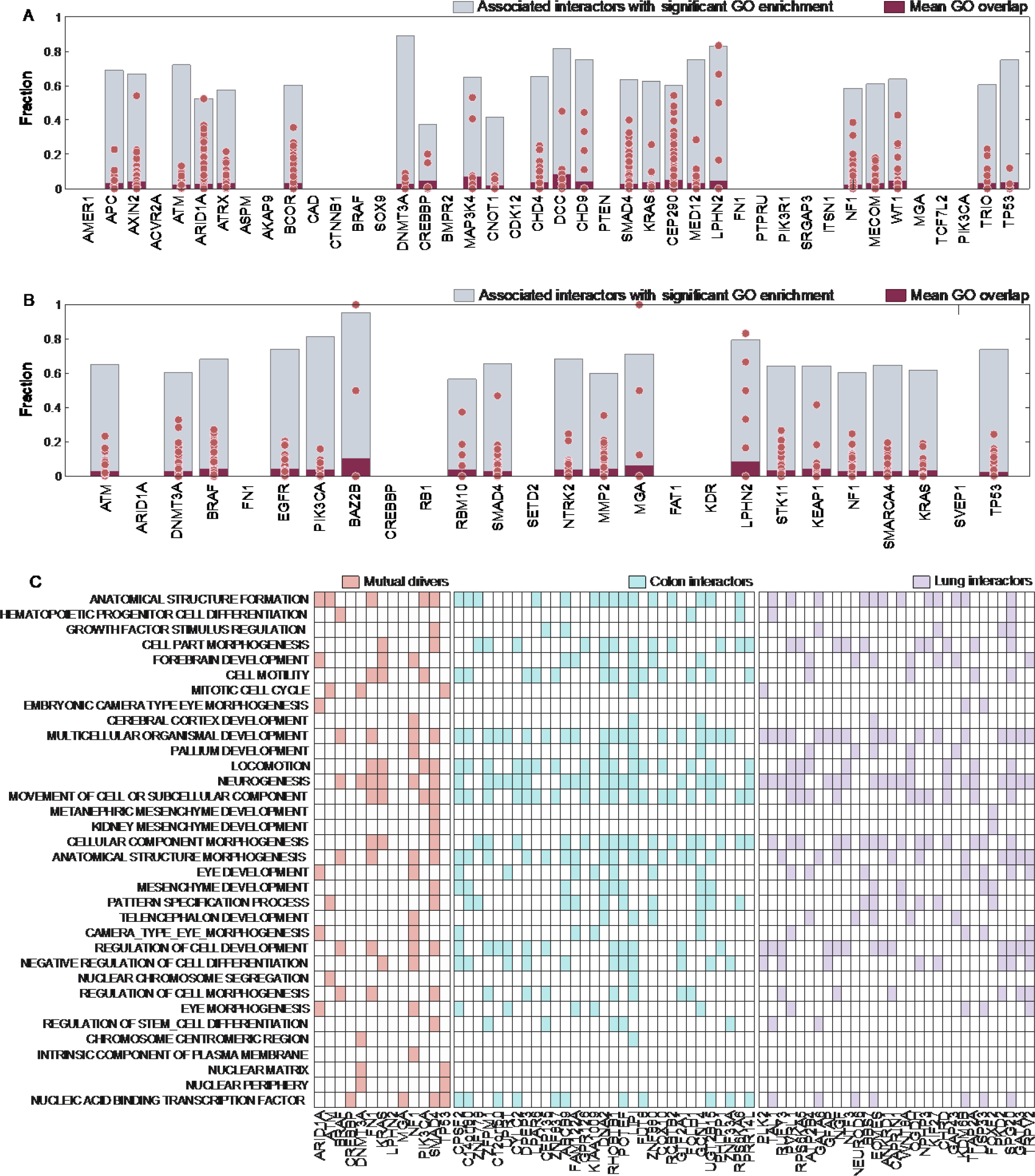
GO enrichment of the predicted interactions of major cancer drivers. (A)-(B) The grey bars show the fraction of GO processes associated with each major cancer driver that are significantly shared with its predicted interactors, for colon and lung cancers, respectively. The dot plots show the percentage of overlap of GO processes between each major driver and its predicted interactors (the red bar shows the mean of this distribution). (C) Heatmaps for GO processes enriched with the shared colon and lung major drivers and their interactors (presented are the interactors that are most strongly associated with these GO processes; for full information, see SI Appendix, Table S6).

For the 13 major drivers that are shared between colon and lung cancers, we identified 1119 GO significantly enriched processes (enrichment P-value < 0.01, SI Appendix, Table S6). For the predicted interactors of these major drivers, 240 GO processes are highly enriched in the case of colon cancer, and 229 processes are highly enriched for lung cancer. Of these, 133 GO processes are shared between the interactors from colon and lung cancers (hyper-geometric P-value ∼=0), and among them, 34 GO processes are shared with the set of processes enriched among drivers (hyper-geometric P-value = 0.02, Fig. 4C, SI Appendix, Table S6). Among the GO processes that are shared between these major cancer drivers and their interactors, are cell motility pathways, regulation of cell development, differentiation and growth factor stimulation, and mesenchyme development processes. It appears plausible that the predicted interactions of diverse genes with the major drivers promote tumor growth and aggressiveness through the modulation of those processes.

### Driver-interactor modules are associated with patients’ survival

We next investigated the bipartite network of major drivers and their predicted interactors. In an attempt to infer the contributions of these interactions to tumor fitness through patients’ survival, we first sought to identify modules of major drivers that share interactors. To this end, we performed a heuristic search for driver modules, aiming to cover the maximum number of drivers in each tumor type. Specifically, we searched for the maximum partition of each tumor graph into disjoint sub-graphs, such that each sub-graph is complete (more precisely, the relation between the modules of drivers and interactors are complete; see Methods for details). In colon cancer, we identified three mutually exclusive modules of drivers (with mutually exclusive pairwise interactions) which together cover 22 of the 23 major colon cancer drivers with predicted interactions (all but the DCC gene), and 47 interactors (Fig. 5A-C). For each module, we then identified the TCGA colon samples in which the given module is highly mutated (see Methods for details). We performed Kaplan Meier survival analysis comparing the survival curves between samples with high vs. low number of mutations of the predicted interactors within each module, conditional on the drivers in the respective module are highly mutated. Strikingly, we found that the high mutation rate of interactors of each of the three modules is associated with poor survival when the driver component of the module is mutated (Fig. 5D-F). For some of these shared interactors, we also find that individual mutations are significantly associated with lower survival rate in the context where their driver module is highly mutated (Fig. 5G-I). Among these, SIX4 expression has been shown to correlate with lymph node metastasis, late stage and unfavorable prognosis of colorectal cancer (51), and PBX3 mutations have been identified in colorectal tumor cells undergoing epithelial-mesenchymal transition, and have been shown to be associated with poor prognosis (52).

**Fig. 5.**
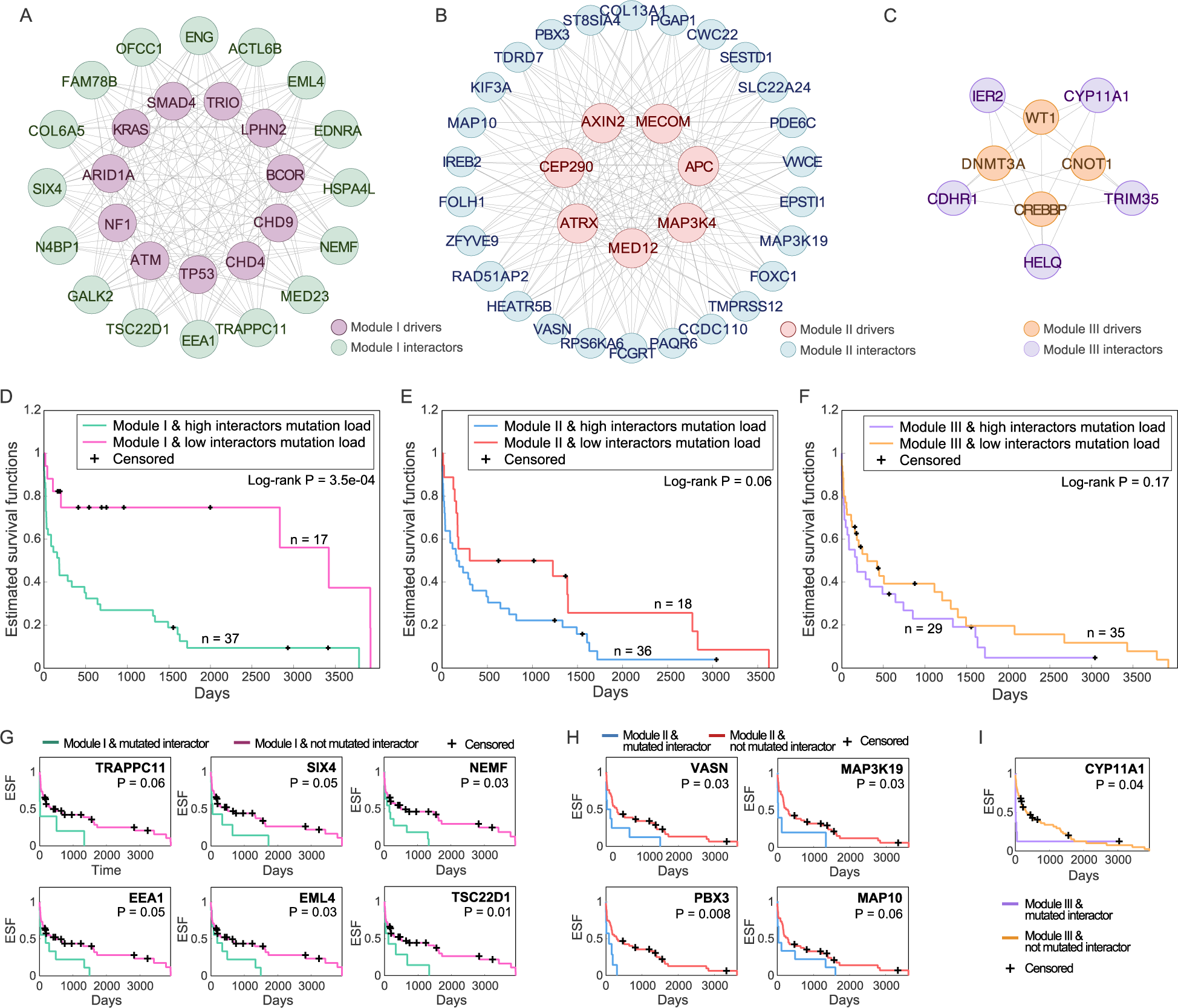
Modules of drivers and interactors in colon cancer. (A) –(C) The complete networks of interactions between modules of major colon cancer drivers (module I-III, respectively), and their predicted shared interactors. (D)-(F) Kaplan Meier survival curves of TCGA colon cancer samples with high mutation rate of drivers modules I-III, respectively, with high vs. low number of mutation in the interactors of these modules (defined by the median). (G)-(I) Kaplan Meier survival curves of TCGA colon cancer samples with vs. without mutations in individual interactors of these modules

For the lung adenocarcinoma mutational data, we detected two mutually exclusive modules of drivers (with mutually exclusive pairwise interactions) that together cover 10 of the 18 major lung cancer drivers with predicted interactions, with 47 interactors of these drivers (Fig. 6A-B). Similarly to the observations on colon cancer, a high number of mutations in the interactors of each driver module is associated with poor survival in samples where the corresponding drivers are mutated (Fig. 6C,E) and with lower progression-free survival (Fig. D,F). For four of the predicted interactors of module 1 (but none for module 2), we also find that some of the individual interacting mutations are significantly associated with lower survival rate when drivers from this module are mutated (Fig. 6G). One of such interactors is EPN1, an Epsin family member shown to regulate tumors progression (53).

**Fig. 6.**
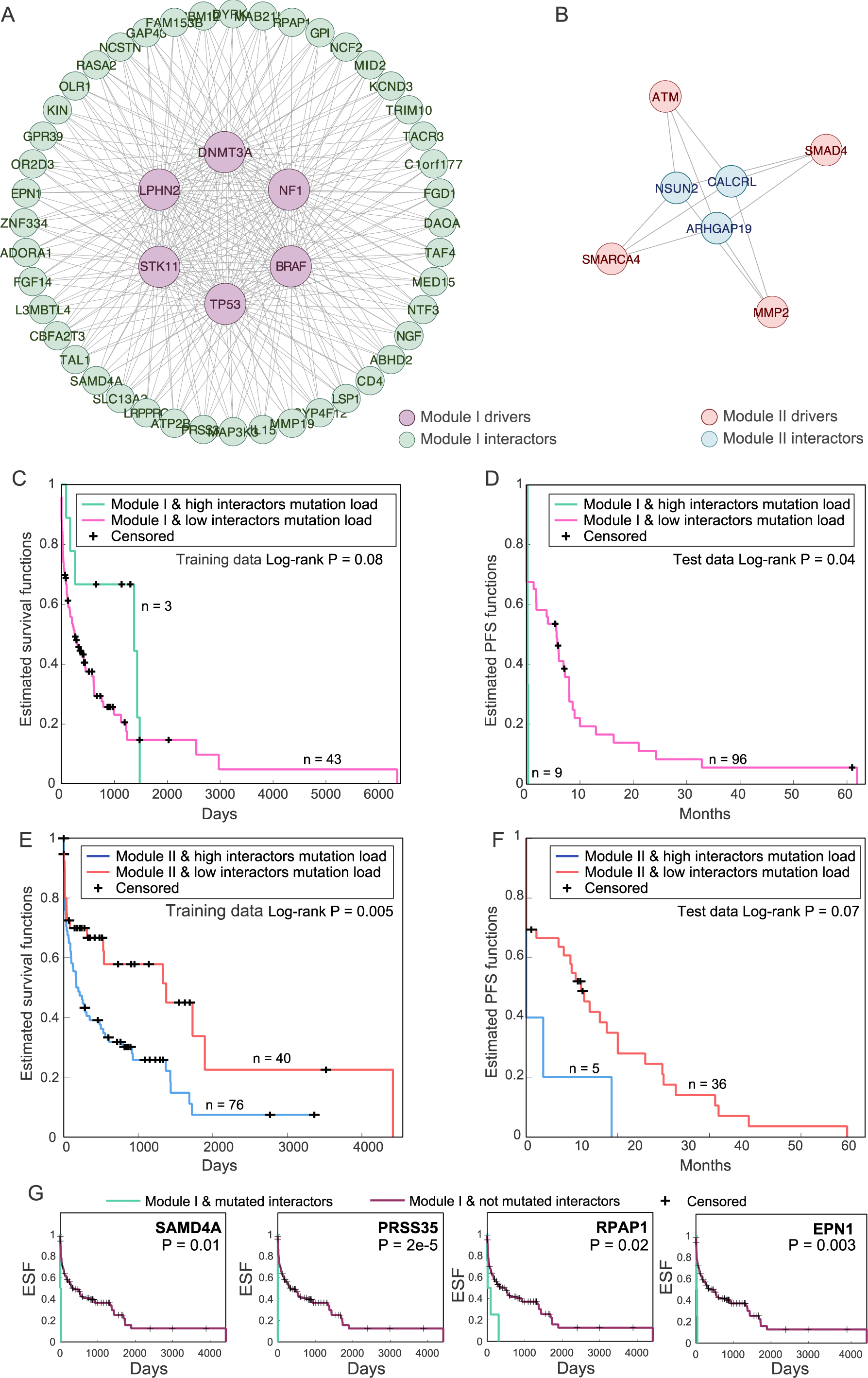
Modules of drivers and interactors in lung cancer. (A)–(B) The complete networks of interactions between modules of major lung cancer drivers (module 1 and 2, respectively) and their predicted shared interactors. (C)-(F) Kaplan Meier survival curves of TCGA colon cancer samples (overall survival, C,E) and the lung cancer test set samples (progression free survival (PFS), D,F) of samples with high mutation rate of driver modules I and II, respectively, with high vs. low number of mutation in the interactors of these modules (defined by the median). (G) Kaplan Meier survival curves of TCGA colon cancer samples with high mutation rate of driver module I (defined by the median), with vs. without mutations of individual interactors of module I.

## Discussion

Most epithelial cancers are preceded by pre-malignant lesions which frequently display mutations in cancer driver genes (54–57). Nevertheless, early and late events are not universally characterized in most epithelial tumors, with the exception of colorectal cancer, where tumor progression has been thoroughly dissected in terms of stepwise accumulation of somatic mutations. This phenomenon starts from transformation of normal epithelium to an adenoma, proceeding to *in situ* carcinoma, and ultimately to invasive and metastatic tumor, where specific mutations mark each step in this tumorigenic transformation (17,19). It is hence reasonable to surmise that time series prediction approaches could be valuable when applied to the sequence of these genetic events in tumors.

Recurrent neural networks (RNNs), and particularly, gated RNN architectures such as LSTMs, have recently shown promising results in learning long-term dependencies of sequences for multiple tasks of classification (58–60), and for data labeling and synthesis (24,61). Here we show that the mutational time series could be utilized via LSTM networks to achieve good performance in otherwise difficult prediction tasks. Using the estimated order of mutations appearance in tumor evolution, we demonstrate that end point conditions, such as the mutational burden and clinical phenotypes, could be predicted from a limited number of mutations. The non-linear relations learned by the networks, together with the discrete representation of the data enable performance, that is significantly superior to the previous models built for this task (62,63). The model can learn intricate dynamics of the mutational sequence in tumor evolution, that can subsequently be used for the reconstruction of mutation sequences. When more data becomes available and the relevant neural network models are further refined, similar approaches could be applied to reconstruct data on actual DNA sequences and could thus extensively contribute to our understanding of tumor evolution. It is worth pointing out that applications of LSTMs generally take advantage of much larger data for training because in these, the input alphabet as well as possible labels are from a much larger range, thus substantially increasing the size of the data set required for training. In our analysis, discretizing the mutational data and maintaining discrete labels (i.e. both input size and labels are always of size 2) generates a simple enough problem that can be approached even with limited amounts of data (Table 1). We also found that LSTM perform much better than linear classifiers, such as Support Vector Machine (SVM), for the prediction of the mutational load. This is likely to be the case because LSTM learn non-linear relationships between mutations in the given sequence, rather than defining a linear separating hyperplane, with the underlining assumption that the mutational load can be predicted using a linear function of mutations.

A potentially important contribution of this approach is the identification of interactors of the major cancer drivers. Here we predict many such interactors and show that they are significantly enriched with STRING interactions and show non-random overlap of GO processes with the corresponding major drivers. Anecdotally, at least some of the better characterized interactors were found to be involved in the same pathway with the corresponding driver. In contrast, we find that the predicted interactors of each drivers are not located on similar chromosomal arms (SI Appendix, Fig. S8), suggesting that these are mostly functional, rather than physical interactions. Furthermore, and most strikingly, we found that mutations in these predicted interactors, in the presence of the corresponding driver mutations, are associated with poor survival, and thus, are likely to confer growth advantage at the respective steps of tumor progression. We identified unique modules of major drivers and their interactors for both colon and lung cancer. The strong correlation with patients’ survival suggests that these interactions are clinically relevant and if further tested, could potentially be used for patients’ stratification and clinical decision-making. The identified interactors can be regarded as secondary drivers whose oncogenic activity is conditional to and associated with the occurrence of mutations in major drivers for the respective cancer types. Hence these conditional drivers could be readily predicted notwithstanding the low accuracy of prediction for most of the major drivers themselves.

To summarize, in this work, we present the application of LSTM for learning the stepwise sequence of mutations in tumors. This approach is shown to efficiently tackle several tasks that are not amenable to standard techniques, such as prediction of the occurrence of mutations and reconstruction of mutational data. Our findings reveal a unidirectional relation between driver and passenger mutations: drivers determine the course of tumorigenesis and their occurrence is difficult to predict from the rest of the mutational landscape, whereas passengers are often linked to specific drivers and so can be predicted with confidence. Thus, drivers are indeed in the driver’s seat and bring with them a host of associated “passengers”, some of which could be secondary, conditional drivers. The present results support the notion that long-term dependencies between genes involved in tumorigenesis and cancer progression are widespread in tumor evolution and can be learned from the mutational sequence using LSTM networks and similar approaches. We show that this notion holds for colon cancer, where the stepwise process of mutation acquisition is established, but also for lung cancer for which such a stepwise model has been suggested but remains controversial. Similar strategies could be readily employed for other tumor types and different types of biological data to advance our understanding of tumor initiation and progression through the dissection of the sequence of evolutionary events.

## Acknowledgments

We thank Koonin group members for helpful discussions. The authors’ research is supported by the Intramural Research Program of the National Institutes of Health (National Library of Medicine).

## Author Contributions

NA conceived of the project; NA, YIW and EVK designed the study; NA performed research; NA and EVK analyzed the results; NA and EVK wrote the manuscript that was read, edited and approved by all authors.

## Methods

### Datasets

Mutational data of colon and lung adenocarcinoma from TCGA were used for training throughout this work and were obtained from Xena browser (64) http://xena.ucsc.edu/. Additional datasets used for testing were obtained from cBioPortal (65,66). The datasets are summarized in Table 1, spanning 1626 samples overall, each derived from a distinct tumor.

### Driver mutation lists for lung and colon cancers

The driver mutations for each tumor type were obtained from (70) and https://www.intogen.org (71). The union of the two lists comprised a comprehensive list of cancer drivers for lung and colon cancers (SI Appendix, Table S7).

### Pre-processing and sorting of the datasets

To generate the binary sequence of mutation, we first discretize the mutational data such that each sample *i* is assigned a value of 1 for gene *g* if it has a non-synonymous mutation in *g*. For each cancer type, we then consider all genes that are mutated at least once in all datasets used (training and test datasets), resulting in the totals of 12,322 mutated genes for colon cancer and 12,327 mutated genes for lung cancer.

To sort the mutations of colon and lung adenocarcinoma by their estimated temporal order, we evaluate the following function for each TCGA training dataset:

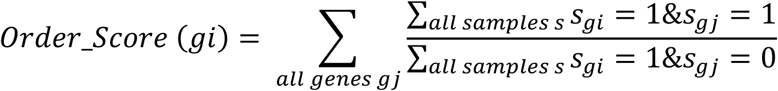

where *sg*_*i*_ = 1 if sample *s* has a mutation in gene *gi*.

We calculate the order score for each considered gene *gi* using the TCGA COAD and LUAD datasets that are used for training. Then, we sort the genes in each dataset by the score assigned to them using the training data of the corresponding cancer type. Order scores calculated using the test datasets were found to significantly correlate with that derived from the training set for both colon and lung cancers (SI Appendix, Fig. S9).

### Long Short-Term Memory (LSTM) machines

Long Short-Term Memory (LSTM) networks are a type of Recurrent Neural Networks that use a time series as input for various prediction tasks. Each LSTM network unit defines a time *t* in a sequence (the subscript *t* denotes the mutation that is ordered *t* in the tumor evolution via the order score defined above), and is composed of the following components:

1. *f*_*t*_ = *σ*_*g*_(*W*_*f*_*x*_*t*_ + *U*_*f*_*h*_*t*-*1*_ + *b*_*f*_)
2. *i*_*t*_ = *σ*_*g*_(*W*_*i*_*x*_*t*_ + *U*_*i*_*h*_*t*-*1*_ + *b*_*i*_)
3. *o*_*t*_ = *σ*_*g*_(*W*_*o*_*x*_*t*_ + *U*_*o*_*h*_*t*-*1*_ + *b*_*o*_)
4. *c*_*t*_ = *f*_*t*_⊙*c*_*t*-*1*_ + *i*_*t*_⊙*σ*_*c*_(*W*_*f*_*x*_*t*_ + *U*_*f*_*h*_*t*-*1*_ + *b*_*f*_)
5. *h*_*t*_ = *f*_*t*_⊙*σ*_*c*_(*c*_*t*_)

where the initial values are *c*_0_ = 0 and *h*_0_ = 0. ⊙ denotes the Hadamard product.

*x*_*t*_ are the input vectors to the LSTM unit (an ordered sequence of mutations). *f*_*t*_, *i*_*t*_ and *o*_*t*_ are the activation vectors for the forget gate, input gate and output gate, respectively. *h*_*t*_ is the output vector of the LSTM unit, and *c*_*t*_ is cell state vector. *W* and *U* are the weight matrices and *b* are the bias matrices that are learned during training. *σ* are the non-linear functions, where *σ*_*g*_ is the gate activation, sigmoid function:

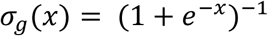

and *σ*_*c*_ is the state activation, *tanh* (hyperbolic tangent) function:

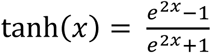

All LSTM networks used in this work are sequence-to-label LSTMs with 5 hidden layers, were trained using Adam optimizer (72), where the maximum number of epochs for training was set to 100 for the mutational load prediction, and to 10 for all other prediction tasks. These thresholds were selected for the sake of computational feasibility; increasing those is expected to further improve the performance. The mini-batch size used for each training iteration was set to 27, with a standard gradient-clipping threshold set to 1.

### Training LSTMs to predict the mutational load

To predict the overall mutational burden and evaluate the number of mutations in the sequence that are required for this task, we train LSTMs for the sequence up to each timepoint when starting once from the mutation ordered last (going backwards in tumor evolution) and second from the mutation ordered first in the sequence. The 2-categorial labels *L*_*s*_*∈* {0,1} are defining ‘low’ vs ‘high’ mutational load, where a sample with ‘low’ mutational load is assigned a label of 0 (if the mutational load of a sample is lower than the median of within a dataset) and ‘high’ mutational load is assigned a label of 1 (samples with mutational load higher than the median). For each time point *t*, a *LSTM*_*t*_ is trained on the training set using the sequence up *t* (either from the first or the last ordered mutation) and is then applied to the test set to predict the mutational load using the time sequence of mutations up-to *t*.

For each time point *t*, the resulting scores (obtained via applying *LSTM*_*t*_ on the sequence of latest *t* mutations in the test) are used to evaluate the performance via two measures:

1. The *AUC*_*t*_ resulting from a ROC curve, predicting the ‘low’ vs. ‘high’ categories in the test.
2. The Spearman rank correlation coefficient *ρ*_*t*_ between the *LSTM*_*t*_ scores that are assigned to each test sample (denoting a likelihood of it having ‘high’ mutational load) and the actual mutational load of the sample.

### Training KNN, SVM and logistic regression classifiers to predict the mutational load

KNN (with K=5), SVM (using linear kernel) and logistic regression classifiers were trained on the training sets using: (1) sequences of the latest ordered mutations that were used as input to the LSTMs (up to 50 latest mutations) and (2) sequences of 50 randomly selected mutations. These were trained to predict ‘low’ vs ‘high’ mutational load as described for the LSTMs. They were then tested on the test sets, and the resulting classification scores were correlated with the true mutational load via Spearman rank-correlation.

### Training LSTMs to predict the occurrence of succeeding mutations in the time sequence

To predict the occurrence a mutation ordered *t* in the sequence we train LSTM for the sequence starting from the mutation ordered last up-to timepoint *t* + 1. The 2-categorial labels *L*_*s*_ ∈ {0,1} are defining the occurrence of the mutation. The LSTMs were trained to predict the occurrence of a mutation in each time point *t* using the training sets of colon and lung, and test their performance for predicting the occurrence of these mutations in the test sets.

### Training LSTM networks to simulate mutational data

To synthesize mutational data, we use the full mutational data for each cancer type (i.e. the union of training and test sets for each tumor type). We reconstruct 100 simulated mutational samples for each tumor type, one mutation at a step, from the last-ordered mutation to the first;

For the mutation ordered last, we randomly assign 1 to *freq*_*l*_ of the 100 generated samples, where *freq*_*l*_ is the overall frequency (in percent) of this last mutation.

To assign the occurrence of every other mutation *t* to the reconstructed samples, we train *LSTM*_*t*_ to predict the occurrence of *t* from the sequence starting from the mutation ordered last up-to timepoint *t* + 1, using the all datasets of each tumor type. We then apply *LSTM*_*t*_ to the simulated sequence (that has been synthesized up-to time point *t* + 1), to obtain a vector of scores predicting the occurrence of mutation *t* in each reconstructed sample. We then use *freq*_*t*_, the frequency of the *t* ordered mutation in the genuine datasets, and assign 1 to the *freq*_*t*_ mutations that were assigned with highest scores by *LSTM*_*t*_ when applied to the reconstructed sequences. PCA analysis was applied to the integration of all datasets (Figures 2D-E, SI Appendix, Fig. S5A-D) or, when inferring the PCA coefficient, without the training sets (SI Appendix, Fig. S5E-F).

### Training LSTMs to identify mutations that interact with the major cancer drivers

To learn the association of mutations with the major driver genes in colon and lung adenocarcinomas, we select the driver genes in which mutations are observed frequently in out training sets, within the top 0.1 percentile. These genes are defined as major drivers and are used as an ordered sequence of mutations (n=42 for colon and n=26 for lung, SI Appendix, Table S4). The sequence of occurrences of these major drivers is used to predict the occurrence of other mutations, excluding those with very low frequency (genes that are mutated in 3 samples or less in the training data are excluded), as the prediction of those could be obtained easily by chance. To predict the occurrence of each mutation, we trained 42 LSTMs for colon and 26 for lung (using the sequence of major drivers up to each time point). The genes that could be predicted with AUC>0.85 for the test set (using the average AUC for the two test sets of colon) repeatedly, from multiple locations in the sequence of drivers (higher than the average number of locations with good performance), are selected as predicted interactors of the major drivers. A graphical schema describing the steps of this analysis can be found in SI Appendix, Fig. S10.

### Assigning major drivers to interacting mutations

To investigate which of the major drivers contribute to the prediction of mutations in each driver-interacting gene (and are hence can be predicted to interact with it), we use the scores produced by the LSTMs where the driver-interacting gene is well-predicted. We then correlate these scores with the occurrence of each of the major drivers, and the major drivers whose occurrence is significantly correlated (Spearman correlation P-value<0.05) with the LSTM scores predicting a given driver-interacting gene are combined with it into driver-interactor pairs.

### Enrichment with STRING interactions and GO analysis

To investigate whether the pairs of major drivers and their predicted interactors are enriched with established interactions, we performed the following analyses:

1. STRING enrichment. Hyper-geometric enrichment analysis was performed for each major driver gene, to find if its LSTM-predicted interactors are enriched with its interactors from the STRING database.
2. GO enrichment. For each major driver, we calculated the percentage of its associated interactors that share a significant number of GO processes with it (hyper-geometric P-value <0.05) and the mean percentage of overlapping GO pathways with its interactors.

We then calculate an empirical P-value from produced via 1000 repetitions, with drivers randomly assigned to the identified interacting mutations (and same degree, i.e. the number of predicted interactions, preserved for each driver), in order to evaluate the probability of obtaining results with a similar or higher significance level by chance.

### Modules of major drivers and interacting genes

We aim to identify modules of major drivers and their interactors, such that the modules are mutually exclusive in terms of both drivers and interactors (i.e. no shared drivers or interactors between the modules). To cover as many drivers as possible, we created modules via a heuristic search using 10,000 repetitions of the following genetic algorithm. For each repetition, we start with a randomly selected major driver. Then, in each round, we randomly select a major driver that has not yet been added to the module. We then add it to the module if its addition does not decrease the number of mutual module-interactors by more than 20%. The round ended when 100 random selections were not added to the current module or when the number of interactors of a module is less than 3. After 10,000 repetitions, we investigated the resulting 10,000 modules and selected those that together cover maximal number of major drivers such that the modules were mutually exclusive with respect to both drivers and interactors.

## Statistical analyses

1. Boxplots and comparisons. For all boxplots, center lines indicate medians, box edges represent the interquartile range, whiskers extend to the most extreme data points not considered outliers, and the outliers are plotted individually using the ‘+’ symbol. Points are defined as outliers if they are greater than *q*_3_+ *w* × (*q*_3_– *q*_1_) or less than *q*_1_– *w* × (*q*_3_– *q*_1_), where *w* is the maximum whisker length, and *q*_1_and *q*_3_are the 25th and 75th percentiles of the sample data, respectively. All differential expression and distribution comparisons P-values are obtained via one-sided Rank-sum test.
2. Survival analyses. All Kaplan Meier analyses are performed by comparing the survival of patients with high scores (module mutational count larger than median) to those with low scores (module mutational count smaller or equal to the median) using a one-sided log-rank test.
3. Correlation coefficients. All correlations coefficients and P-values were obtained using the Spearman rank correlation test.
4. Enrichment analysis. All enrichment P-values were calculated using the hypergeometric enrichment test.
5. Distribution comparisons. All comparisons between two distributions were performed using the one-sided Rank-sum test.

### Code availability

All code was implemented in MATLAB_R2018a using Deep Learning Toolbox and is publicly available in GitHub: https://github.com/noamaus/LSTM-Mutational-series

## References

1. Vogelstein B, Kinzler KW. The multistep nature of cancer. Trends in Genetics. 1993. p. 138–41.

2. Farber E. The Multistep Nature of Cancer Development. Cancer Res. 1984;44(10):4217– 23.

3. Stratton MR, Campbell PJ, Futreal PA. The cancer genome. Nature. 2009. p. 719–24.

4. Vogelstein B, Papadopoulos N, Velculescu VE, Zhou S, Diaz LA, Kinzler KW. Cancer genome landscapes. Science. 2013. p. 1546–58.

5. Pleasance ED, Cheetham RK, Stephens PJ, McBride DJ, Humphray SJ, Greenman CD, et al. A comprehensive catalogue of somatic mutations from a human cancer genome. Nature [Internet]. 2009;463(7278):191–6. Available from: http://www.nature.com/doifinder/10.1038/nature08658.1038/nature08658

6. McFarland CD, Korolev KS, Kryukov G V., Sunyaev SR, Mirny LA. Impact of deleterious passenger mutations on cancer progression. Proc Natl Acad Sci [Internet]. 2013;110(8):2910–5. Available from: http://www.pnas.org/cgi/doi/10.1073/pnas.1213968110

7. Persi E, Wolf YI, Leiserson MDM, Koonin E V., Ruppin E. Criticality in tumor evolution and clinical outcome. Proc Natl Acad Sci [Internet]. 2018;115(47):E11101–10. Available from: http://www.pnas.org/lookup/doi/10.1073/pnas.1807256115

8. Tanaka T. Colorectal carcinogenesis: Review of human and experimental animal studies. J Carcinog [Internet]. 2009;8(1):5. Available from: http://www.carcinogenesis.com/text.asp?2009/8/1/5/49014

9. Loeb LA, Harris CC. Advances in chemical carcinogenesis: A historical review and prospective. Cancer Research. 2008. p. 6863–72.

10. Knudson AG. Two genetic hits (more or less) to cancer. Nat Rev Cancer [Internet]. 2001;1(2):157–62. Available from: http://www.nature.com/articles/35101031

11. Teixeira MR, Heim S. Multiple numerical chromosome aberrations in cancer: What are their causes and what are their consequences? Seminars in Cancer Biology. 2005. p. 3–12.

12. Gillies RJ, Verduzco D, Gatenby RA. Evolutionary Dynamics Unifies Carcinogenesis and Cancer Therapy. Nat Rev Cancer. 2012;12(7):487–93.

13. Vogelstein B, Fearon ER, Hamilton SR, Kern SE, Preisinger AC, Leppert M, et al. Genetic Alterations during Colorectal-Tumor Development. N Engl J Med [Internet]. 1988;319(9):525–32. Available from: http://www.nejm.org/doi/abs/10.1056/NEJM198809013190901

14. Armaghany T, Wilson JD, Chu Q, Mills G. Genetic alterations in colorectal cancer. Gastrointest Cancer Res [Internet]. 2012;5(1):19–27. Available from: http://www.pubmedcentral.nih.gov/articlerender.fcgi?artid=PMC3348713

15. Fearon ER. Genetic alterations underlying colorectal tumorigenesis. Cancer Surv [Internet]. 1992;12:119–36. Available from: http://www.ncbi.nlm.nih.gov/entrez/query.fcgi?cmd=Retrieve&db=PubMed&dopt=Citation&list_uids=1638544

16. Lurje G, Zhang W, Lenz H-J. Molecular prognostic markers in locally advanced colon cancer. Clin Colorectal Cancer [Internet]. 2007;6(10):683–90. Available from: http://dx.doi.org/10.3816/CCC.2007.n.037

17. Noguchi M. Stepwise progression of pulmonary adenocarcinoma-clinical and molecular implications. Cancer and Metastasis Reviews. 2010. p. 15–21.

18. Sanada Y, Yoshida K, Ohara M, Oeda M, Konishi K, Tsutani Y. Histopathologic evaluation of stepwise progression of pancreatic carcinoma with immunohistochemical analysis of gastric epithelial transcription factor SOX2: Comparison of expression patterns between invasive components and cancerous or nonneoplastic intr. Pancreas. 2006;32(2):164–70.

19. Yatabe Y, Borczuk AC, Powell CA. Do all lung adenocarcinomas follow a stepwise progression? Lung Cancer. 2011. p. 7–11.

20. Hochreiter S, Urgen Schmidhuber J. LONG SHORT-TERM MEMORY. Neural Comput [Internet]. 1997;9(8):1735–80. Available from: http://www7.informatik.tu-muenchen.de/~hochreit%5Cn http://www.idsia.ch/~juergen

21. Williams RJ, Zipser D. A Learning Algorithm for Continually Running Fully Recurrent Neural Networks. Neural Computation. 1989. p. 270–80.

22. Gers FA, Schmidhuber J, Cummins F. Learning to forget: Continual prediction with LSTM. Neural Comput. 2000;12(10):2451–71.

23. Schmidhuber J, Wierstra D, Gomez F. Evolino: Hybrid neuroevolution / optimal linear search for sequence learning. In: IJCAI International Joint Conference on Artificial Intelligence. 2005. p. 853–8.

24. Graves A. Generating sequences with recurrent neural networks. preprint. arXiv:13080850 [Internet]. 2013;1–43. Available from: http://arxiv.org/abs/1308.0850

25. Sundermeyer M, Schlueter R, Ney H. LSTM Neural Networks for Language Modeling. In: Proceedings of INTERSPEECH. 2012. p. 194–7.

26. Lai S, Xu L, Liu K, Zhao J. Recurrent Convolutional Neural Networks for Text Classification. AAAI’15. 2015;2267–73.

27. Sutskever I, Martens J, Hinton G. Generating Text with Recurrent Neural Networks. In: Procededings of the 28th International Conference on Machine Learning (ICML’11) [Internet]. 2011. p. 1017–24. Available from: http://www.icml-2011.org/papers/524_icmlpaper.pdf

28. Weinstein JN, Collisson EA, Mills GB, Shaw KRM, Ozenberger BA, Ellrott K, et al. The Cancer Genome Atlas Pan-Cancer analysis project. Nat Genet [Internet]. 2013;45(10):1113–20. Available from: http://dx.doi.org/10.1038/ng.2764

29. Levine RL, Abdel-Wahab O, Getz G, Mellinghoff IK, Attolini CS-O, Michor F, et al. A mathematical framework to determine the temporal sequence of somatic genetic events in cancer. Proc Natl Acad Sci. 2010;107(41):17604–9.

30. Cheng YK, Beroukhim R, Levine RL, Mellinghoff IK, Holland EC, Michor F. A mathematical methodology for determining the temporal order of pathway alterations arising during gliomagenesis. PLoS Comput Biol. 2012;8(1).

31. Jiang F, Kallioniemi O-P, Schäffer AA, Desper R, Moch H, Papadimitriou CH. Distance-Based Reconstruction of Tree Models for Oncogenesis. J Comput Biol. 2002;7(6):789–803.

32. Hoglund M, Gisselsson D, Mandahl N, Johansson B, Mertens F, Mitelman F, et al. Multivariate analyses of genomic imbalances in solid tumors reveal distinct and converging pathways of karyotypic evolution. Genes Chromosom Cancer. 2001;31(2):156–71.

33. Lin SH, Raju GS, Huff C, Ye Y, Gu J, Chen JS, et al. The somatic mutation landscape of premalignant colorectal adenoma. Gut. 2018;67(7):1299–305.

34. Bamford S, Dawson E, Forbes S, Clements J, Pettett R, Dogan A, et al. The COSMIC (Catalogue of Somatic Mutations in Cancer) database and website. Br J Cancer. 2004;91(2):355–8.

35. Tate JG, Bamford S, Jubb HC, Sondka Z, Beare DM, Bindal N, et al. COSMIC: the Catalogue Of Somatic Mutations In Cancer. Nucleic Acids Res [Internet]. 2018; Available from: https://academic.oup.com/nar/advance-article/doi/10.1093/nar/gky1015/5146192

36. Wolff RK, Hoffman MD, Wolff EC, Herrick JS, Sakoda LC, Samowitz WS, et al. Mutation analysis of adenomas and carcinomas of the colon: Early and late drivers. Genes Chromosom Cancer. 2018;57(7):366–76.

37. Aithal A, Rauth S, Kshirsagar P, Shah A, Lakshmanan I, Junker WM, et al. MUC16 as a novel target for cancer therapy. Expert Opinion on Therapeutic Targets. 2018. p. 675–86.

38. Sur I, Neumann S, Noegel AA. Nesprin-1 role in DNA damage response. Nucl (United States). 2014;5(2).

39. Bozic I, Antal T, Ohtsuki H, Carter H, Kim D, Chen S, et al. Accumulation of driver and passenger mutations during tumor progression. Proc Natl Acad Sci [Internet]. 2010;107(43):18545–50. Available from: http://www.pnas.org/cgi/doi/10.1073/pnas.1010978107

40. Maaten L van der, Hinton G. Visualizing Data using t-SNE. J Mach Learn Res [Internet]. 2008;9:2579–605. Available from: http://www.ncbi.nlm.nih.gov/entrez/query.fcgi?db=pubmed&cmd=Retrieve&dopt=AbstractPlus&list_uids=7911431479148734548related:VOiAgwMNy20J

41. Szklarczyk D, Franceschini A, Wyder S, Forslund K, Heller D, Huerta-Cepas J, et al. STRING v10: Protein-protein interaction networks, integrated over the tree of life. Nucleic Acids Res. 2015;43(D1):D447–52.

42. Szklarczyk D, Morris JH, Cook H, Kuhn M, Wyder S, Simonovic M, et al. The STRING database in 2017: Quality-controlled protein-protein association networks, made broadly accessible. Nucleic Acids Res. 2017;45(D1):D362–8.

43. Zawel L, Le Dai J, Buckhaults P, Zhou S, Kinzler KW, Vogelstein B, et al. Human Smad3 and Smad4 are sequence-specific transcription activators. Mol Cell. 1998;1(4):611–7.

44. Ahn Y-H, Yang Y, Gibbons DL, Creighton CJ, Yang F, Wistuba II, et al. Map2k4 Functions as a Tumor Suppressor in Lung Adenocarcinoma and Inhibits Tumor Cell Invasion by Decreasing Peroxisome Proliferator-Activated Receptor 2 Expression. Mol Cell Biol [Internet]. 2011;31(21):4270–85. Available from: http://mcb.asm.org/cgi/doi/10.1128/MCB.05562-11

45. Baba Y, Nosho K, Shima K, Irahara N, Chan AT, Meyerhardt JA, et al. HIF1A overexpression is associated with poor prognosis in a cohort of 731 colorectal cancers. Am J Pathol. 2010;176(5):2292–301.

46. Papageorgis P, Cheng K, Ozturk S, Gong Y, Lambert AW, Abdolmaleky HM, et al. Smad4 inactivation promotes malignancy and drug resistance of colon cancer. Cancer Res. 2011;71(3):998–1008.

47. Harris CC. p53 tumor suppressor gene: At the crossroads of molecular carcinogenesis, molecular epidemiology, and cancer risk assessment. In: Environmental Health Perspectives. 1996. p. 435–9.

48. Freed-Pastor WA, Prives C. Mutant p53: One name, many proteins. Genes Dev. 2012;26(12):1268–86.

49. Carbon S, Dietze H, Lewis SE, Mungall CJ, Munoz-Torres MC, Basu S, et al. Expansion of the gene ontology knowledgebase and resources: The gene ontology consortium. Nucleic Acids Res. 2017;45(D1):D331–8.

50. Ashburner M, Ball CA, Blake JA, Botstein D, Butler H, Cherry JM, et al. Gene ontology: Tool for the unification of biology. Nature Genetics. 2000. p. 25–9.

51. Li G, Hu F, Luo X, Hu J, Feng Y. SIX4 promotes metastasis via activation of the PI3K-AKT pathway in colorectal cancer. PeerJ [Internet]. 2017;5:e3394. Available from: https://peerj.com/articles/3394

52. Lamprecht S, Kaller M, Schmidt EM, Blaj C, Schiergens TS, Engel J, et al. PBX3 is part of an EMT regulatory network and indicates poor outcome in colorectal cancer. Clin Cancer Res. 2018;24(8).

53. Tessneer KL, Cai X, Pasula S, Dong Y, Liu X, Chang B, et al. Epsin Family of Endocytic Adaptor Proteins as Oncogenic Regulators of Cancer Progression. J Can Res Updates [Internet]. 2013;2:144–50. Available from: http://www.pubmedcentral.nih.gov/articlerender.fcgi?artid=3911794&tool=pmcentrez&rendertype=abstract

54. Ryan BM, Faupel-Badger JM. The hallmarks of premalignant conditions: A molecular basis for cancer prevention. Seminars in Oncology. 2016. p. 22–35.

55. Shin DM, Hong WK, Kim J, Roth JA, Surgery T, Ro JY, et al. Activation of p53 Gene Expression in Premalignant Lesions during Head and Neck Ttomorigenesis. Cancer Res. 1994;54(2):321–6.

56. Viola M V, Fromowitz F, Oravez S, Deb S, Schlom J. ras Oncogene p21 expression is increased in premalignant lesions and high grade bladder carcinoma. J Exp Med [Internet]. 1985;161(5):1213–8. Available from: http://link.springer.com/10.1007/174_2014_1003 http://www.ncbi.nlm.nih.gov/pubmed/3886828%0A http://www.pubmedcentral.nih.gov/articlerender.fcgi?artid=PMC2187613

57. Enomoto T, Tanizawa O, Inoue M. K-ras Activation in Premalignant and Malignant Epithelial Lesions of the Human Uterus. Cancer Res. 1991;51(19):5308–14.

58. Beil J, Perner G, Asfour T. Design and control of the lower limb exoskeleton KIT-EXO-1. IEEE Int Conf Rehabil Robot. 2015;2015–Septe:119–24.

59. Lipton ZC, Berkowitz J, Elkan C. A Critical Review of Recurrent Neural Networks for Sequence Learning. 2015;1–38. Available from: http://arxiv.org/abs/1506.00019

60. Shi X, Chen Z, Wang H, Yeung D-Y, Wong W, Woo W. Convolutional LSTM Network: A Machine Learning Approach for Precipitation Nowcasting. 2015;1–9. Available from: http://arxiv.org/abs/1506.04214

61. Graves A, Fernandez S, Gomez F, Schmidhuber J. Connectionist Temporal Classification: Labelling Unsegmented Sequence Data with Recurrent Neural Networks. In: Proceedings of the 23rd International Conference on Machine Learning (ICML). 2006. p. 369–76.

62. Lyu GY, Yeh YH, Yeh YC, Wang YC. Mutation load estimation model as a predictor of the response to cancer immunotherapy. npj Genomic Med. 2018;3(1).

63. Roszik J, Haydu LE, Hess KR, Oba J, Joon AY, Siroy AE, et al. Novel algorithmic approach predicts tumor mutation load and correlates with immunotherapy clinical outcomes using a defined gene mutation set. BMC Med. 2016;14(1).

64. Goldman M, Craft B, Kamath A, Brooks AN, Zhu J, Haussler D. The UCSC Xena Platform for cancer genomics data visualization and interpretation. bioRxiv [Internet]. 2018;326470. Available from: https://www.biorxiv.org/content/early/2018/05/25/326470

65. Gao J, Aksoy BA, Dogrusoz U, Dresdner G, Gross B, Sumer SO, et al. Integrative analysis of complex cancer genomics and clinical profiles using the cBioPortal. Sci Signal [Internet]. 2013;6(269):pl1. Available from: http://www.ncbi.nlm.nih.gov/pubmed/23550210%5Cnhttp://www.pubmedcentral.nih.gov/articlerender.fcgi?artid=PMC4160307

66. Cerami E, Gao J, Dogrusoz U, Gross BE, Sumer SO, Aksoy BA, et al. The cBio Cancer Genomics Portal: An open platform for exploring multidimensional cancer genomics data. Cancer Discov. 2012;2(5):401–4.

67. Giannakis M, Mu XJ, Shukla SA, Qian ZR, Cohen O, Nishihara R, et al. Genomic Correlates of Immune-Cell Infiltrates in Colorectal Carcinoma. Cell Rep. 2016;15(4):857– 65.

68. Seshagiri S, Stawiski EW, Durinck S, Modrusan Z, Storm EE, Conboy CB, et al. Recurrent R-spondin fusions in colon cancer. Nature. 2012;488(7413):660–4.

69. Imielinski M, Berger AH, Hammerman PS, Hernandez B, Pugh TJ, Hodis E, et al. Mapping the hallmarks of lung adenocarcinoma with massively parallel sequencing. Cell. 2012;150(6):1107–20.

70. Matthew Bailey AH, Tokheim C, Porta-Pardo E, Mills GB, Karchin R, Ding L, et al. Comprehensive Characterization of Cancer Driver Genes and Mutations Article Comprehensive Characterization of Cancer Driver Genes and Mutations. Cell [Internet]. 2018;173:371–376.e18. Available from: https://doi.org/10.1016/j.cell.2018.02.060

71. Rubio-Perez C, Tamborero D, Schroeder MP, Antolín AA, Deu-Pons J, Perez-Llamas C, et al. In Silico Prescription of Anticancer Drugs to Cohorts of 28 Tumor Types Reveals Targeting Opportunities. Cancer Cell. 2015;27(3):382–96.

72. Kingma DP, Ba JL. Adam: a Method for Stochastic Optimization. Int Conf Learn Represent 2015. 2015;1–15.

