## Supplementary material for "*In silico* learning of tumor evolution through mutational time series": SI appendix

**This PDF file includes:**

Figures S1 to S10
Captions for Tables S1 to S7

**Other supplementary materials for this manuscript include the following:**

Table S1 to S7

**Figure S1. (A)** Heatmap showing the order score for colon cancer driver genes. **(B)** Scatter plot showing the correlation between the log-transformed ratio of mutations frequency in colorectal adenomas and that in carcinomas (x-axis) and the order score assigned to each mutation (y-axis). Colon cancer drivers are the red dots, where all other genes are blue.

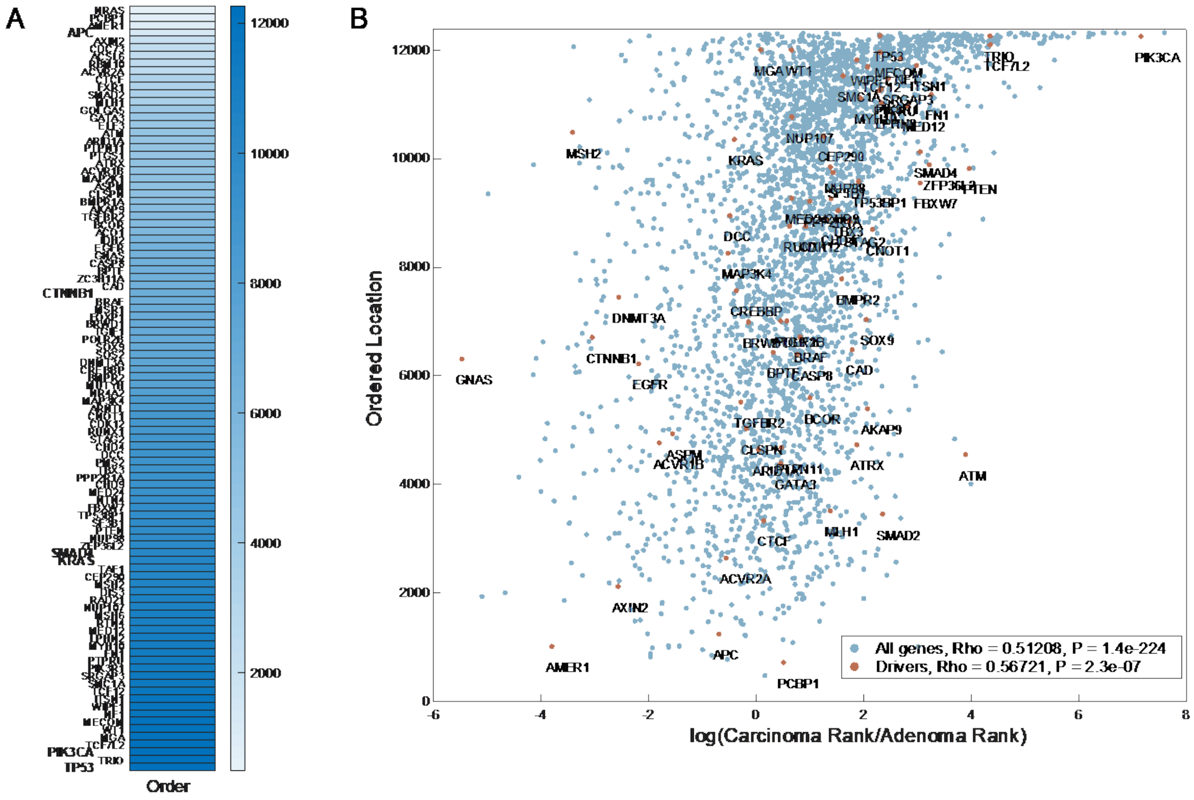

**Figure S2. (A)** The test AUCs (Y-axis) obtained for training LSTMs with different lengths of mutation sequences (X-axis) when starting from the earliest ordered mutation, for colon and lung test sets. **(B)** The rank correlation coefficient (Y-axis) between the true mutational load and the score obtained with training LSTMs with different lengths of mutation sequences (X-axis) when starting from the earliest ordered mutation, for colon and lung test sets.

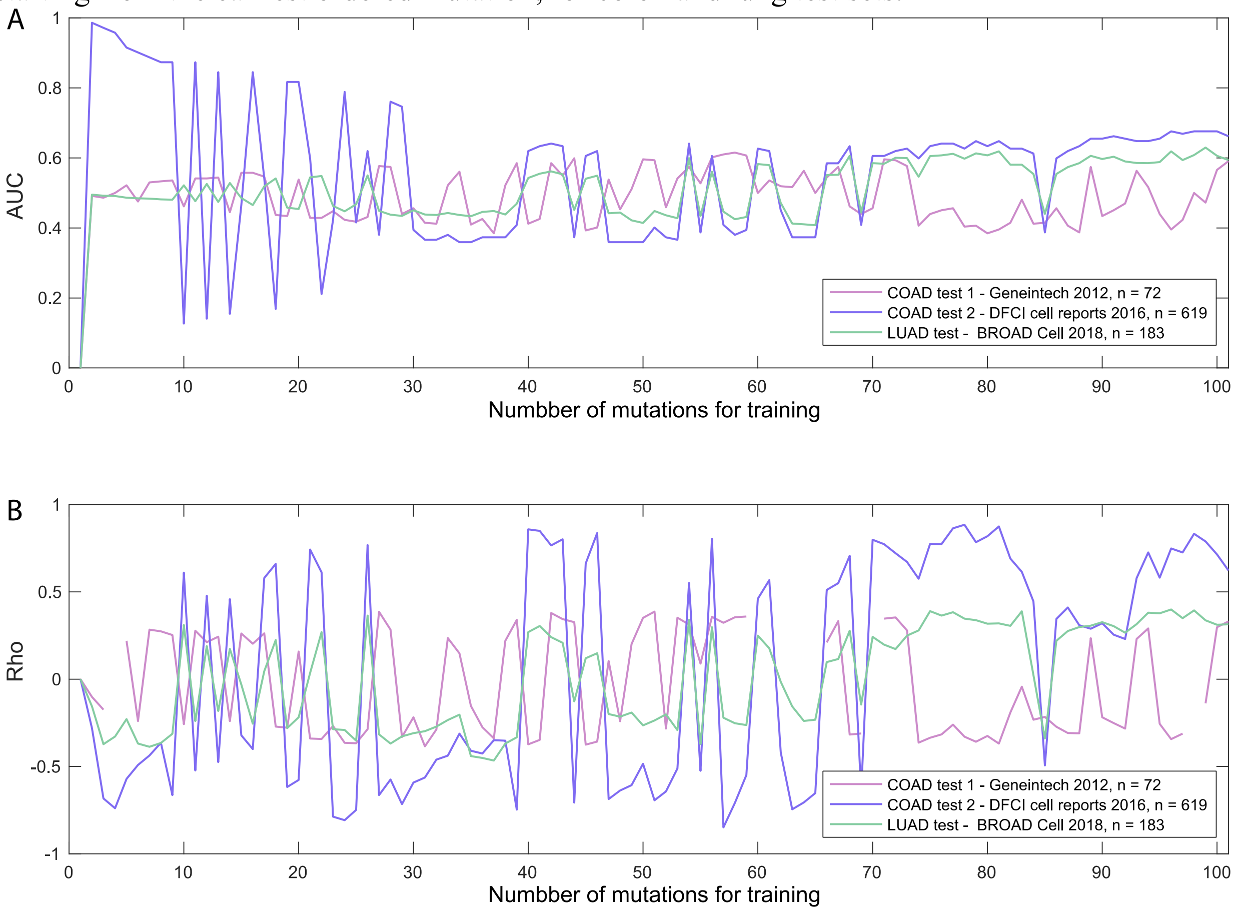

**Figure S3. (A),(C)** Mean AUC of mutation prediction in the sequence for the two colon test sets: comparison of drivers with other genes with mutation frequency in to top 0.5 and 0.75 percentiles, respectively.

**(B),(D)** AUC of mutation prediction in the sequence for the lung test set: comparison of drivers with other genes with mutation frequency in to top 0.5 and 0.75 percentiles, respectively.

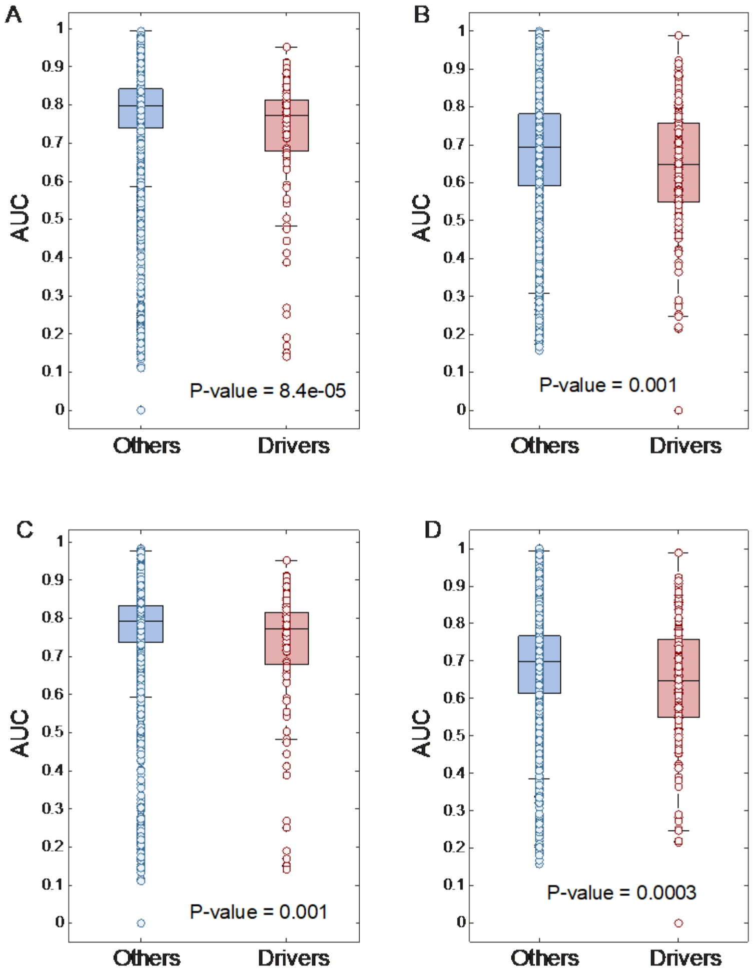

**Figure S4. (A)** Bar plots showing the number of samples, from each original colon cancer dataset and the simulated dataset, that are clustered into each cluster with K-means clustering, for k=2-6. **(B)** Bar plots showing the number of samples, from each original lung cancer dataset and the simulated dataset, that are clustered into each cluster with K-means clustering, for k=2-6. **(C),(D)** scatter plots showing dimension 1-2 obtained from tSNE dimensionality reduction using the combined datasets and the reconstructed data for colon and lung cancer, respectively. The tSNE was applied with Euclidian distance matric, 4 natural clusters of the data, 2 output dimensions, and 30 local neighbors for each point (perplexity).

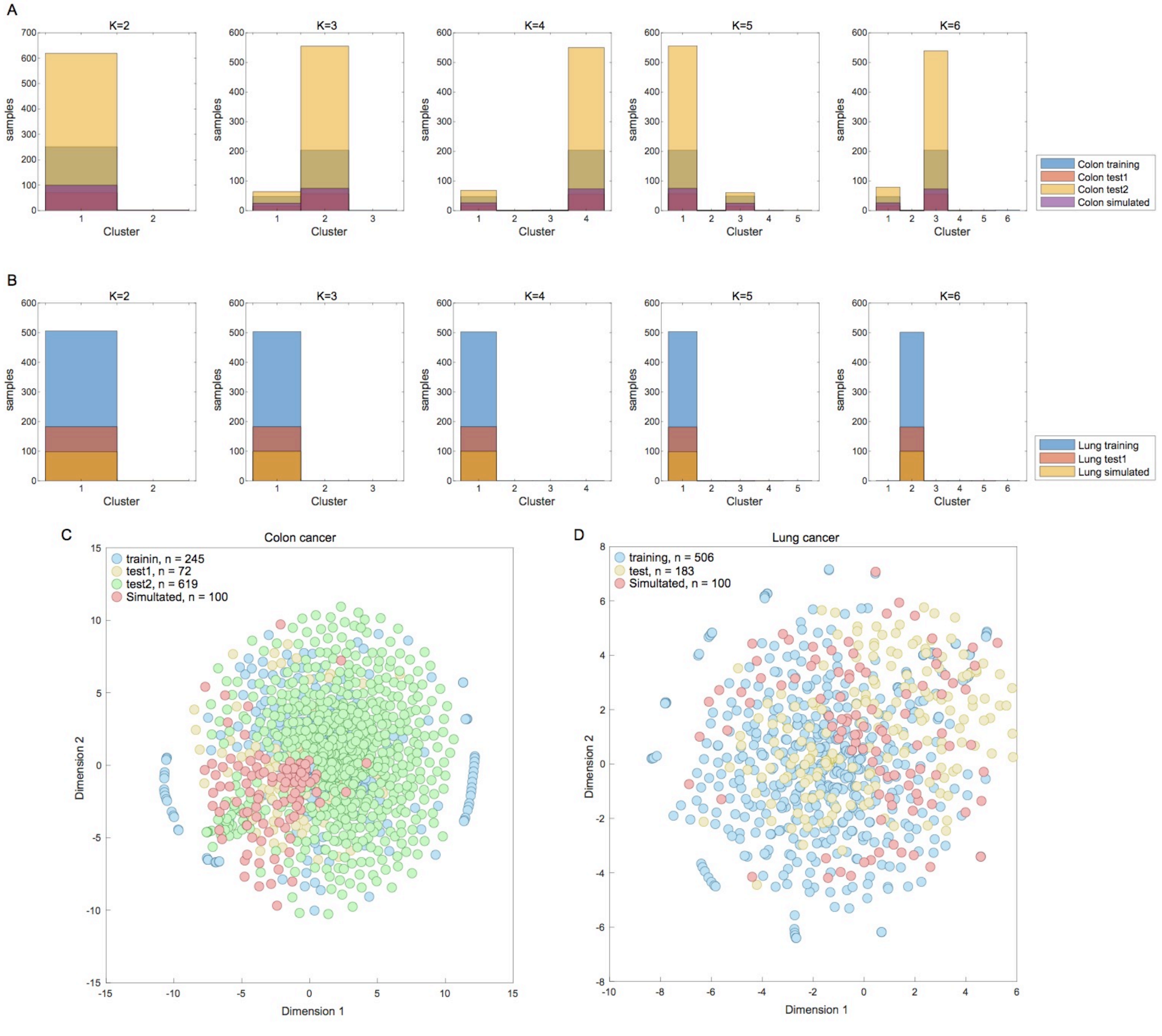

**Figure S5.**

**(A),(C)** Scatter plots of PC1-PC3 obtained by PCA applied to the combined colon mutational data from all datasets used and the colon simulated samples using mutations with the frequency within the top 50%, 99%, respectively. **(B),(D)** Scatter plots of PC1-PC3 obtained by PCA applied to the combined lung mutational data from all datasets used and the lung simulated samples using mutations with the frequency within the top 50%, 99%, respectively. **(E),(F)** **S**catter plots of PC1-PC3 obtained by PCA applied to the combined colon and lung cancer data, respectively, when excluding the training set for obtaining the PCA weights. The parenthesis indicate the percentage of variance explained by each PC.

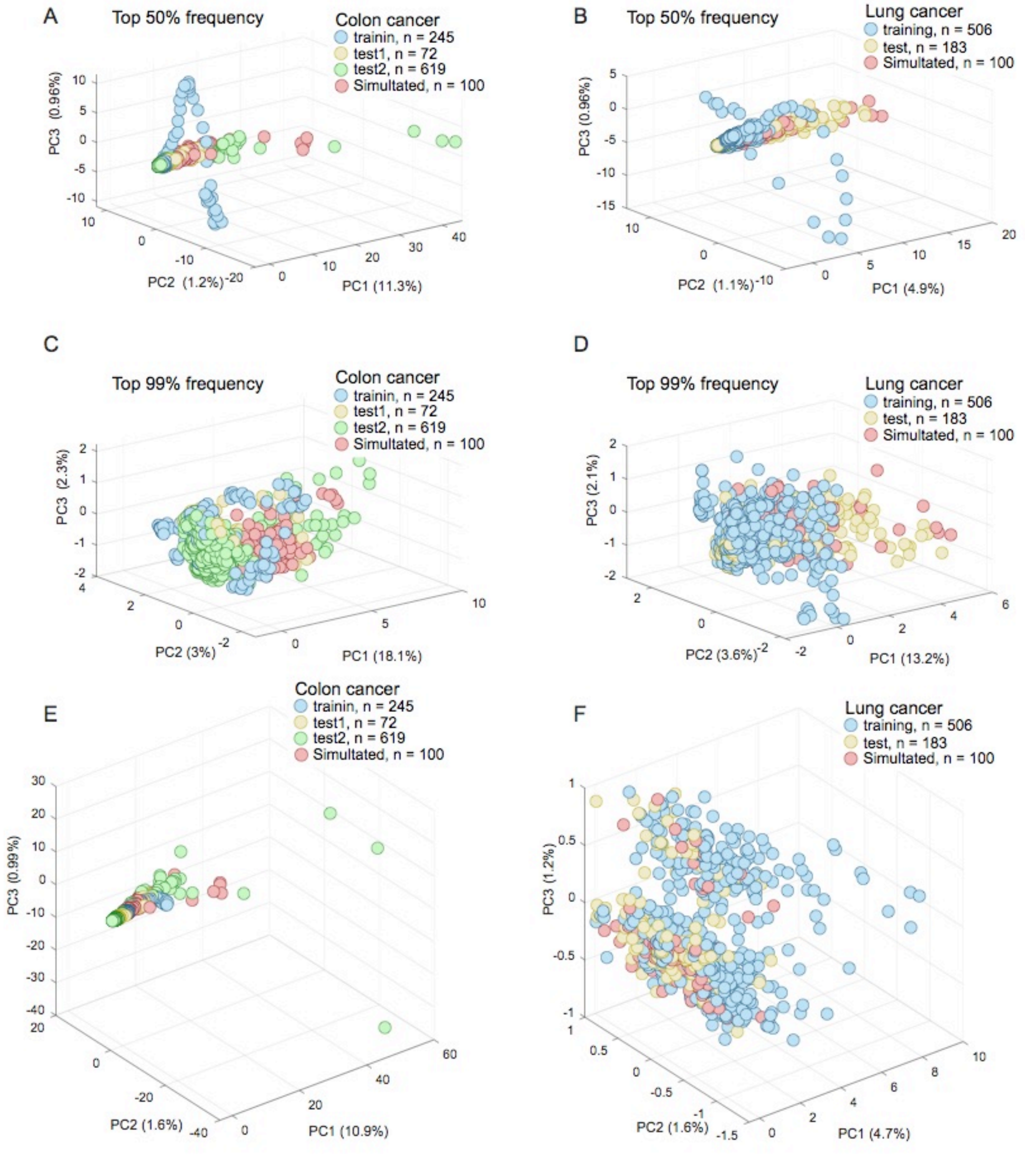

**Figure S6. (A)** Heatmaps showing the high frequency (top 20%) cancer drivers in the colon cancer simulated mutational samples (red), the colon TCGA mutational data (blue) and the colon test sets (yellow, green). **(B)** Heatmaps showing the high frequency (top 20%) cancer drivers in the lung simulated mutational samples (red), the lung TCGA mutational data (blue) and the lung test set (yellow).

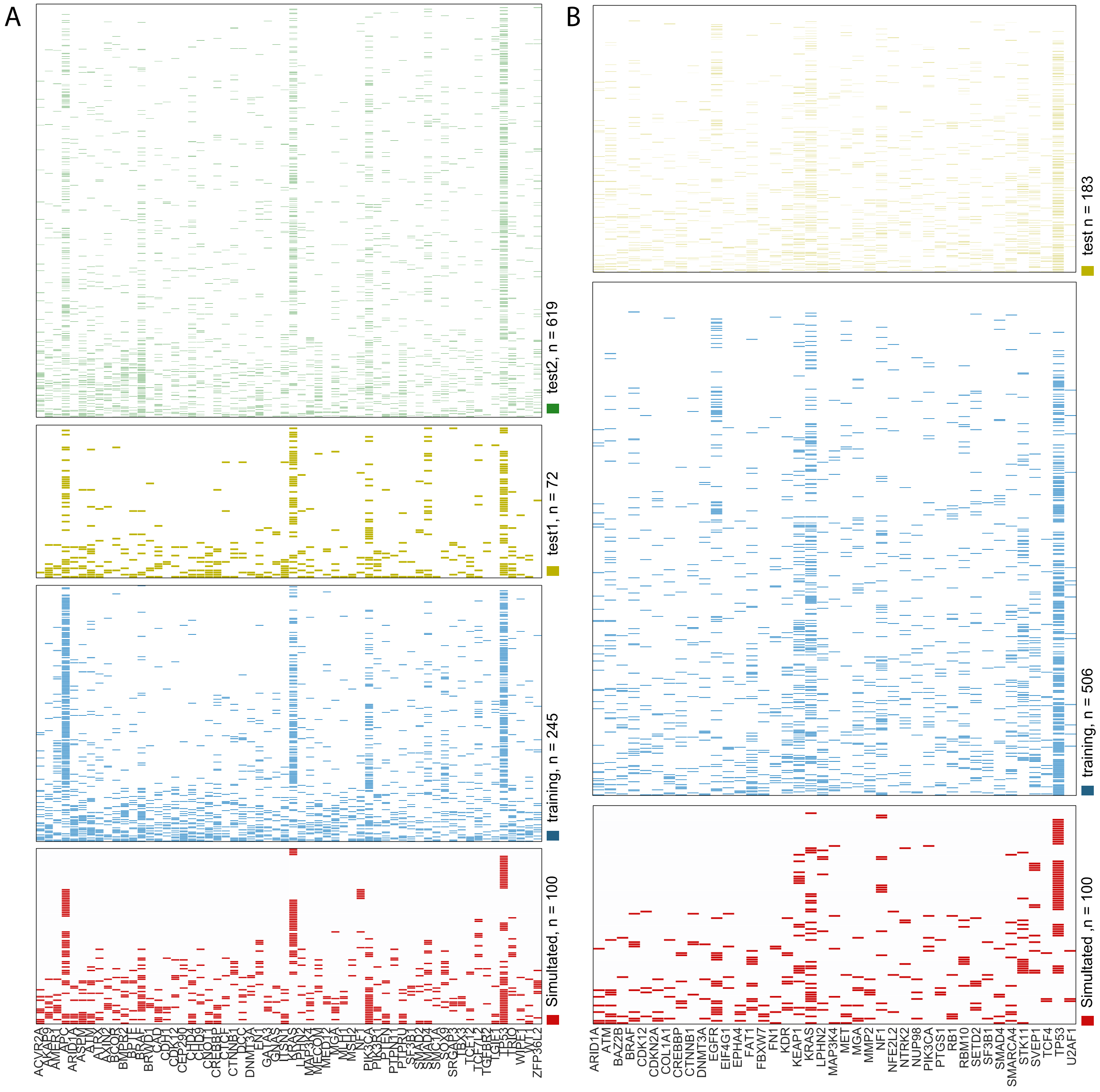

**Figure S7. (A)-(C)** Comparing the prediction performance (AUC) for predicting the occurrence of succeeding mutations in the time sequence, for the last ordered 300 mutations, between the original order function (right panels) vs. 10 shuffled mutation orders (left panels) used for prediction (maintaining the same mutations, but with different random orders), for colon test set 1, colon test set 2 and lung test set, respectively. The P-values are for paired, one-sided rank-sum test, and are indicating the maximal value for all permutations.

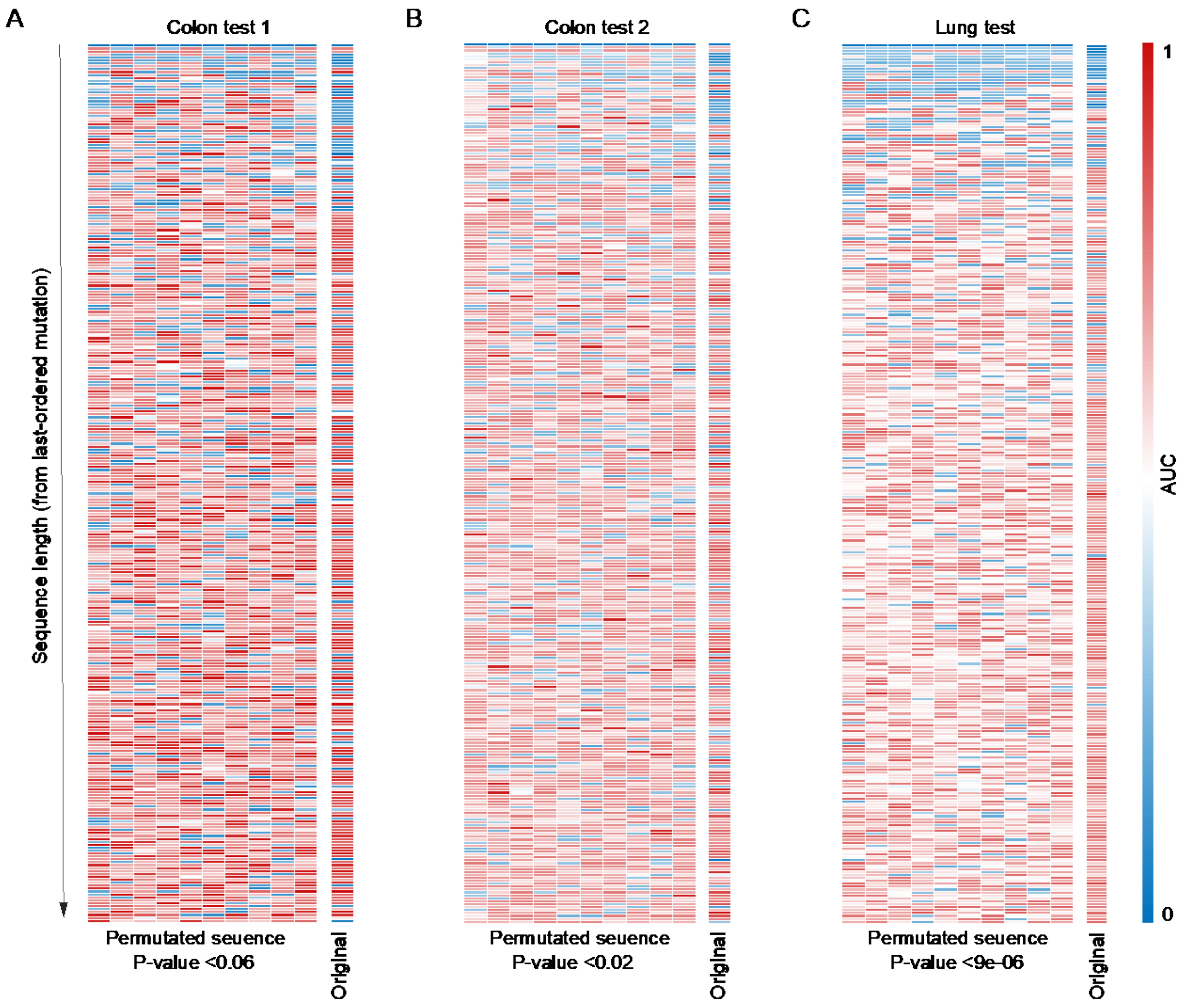

**Figure S8. (A)** Heatmap showing number of predicted interactors for each colon cancer major driver in each of 39 chromosomal arms. The arm to which the driver belong is highlighted in red. **(B)** Heatmap showing number of predicted interactors for each lung cancer major driver in each of 39 chromosomal arms. The arm to which the driver belong is highlighted in red.

A

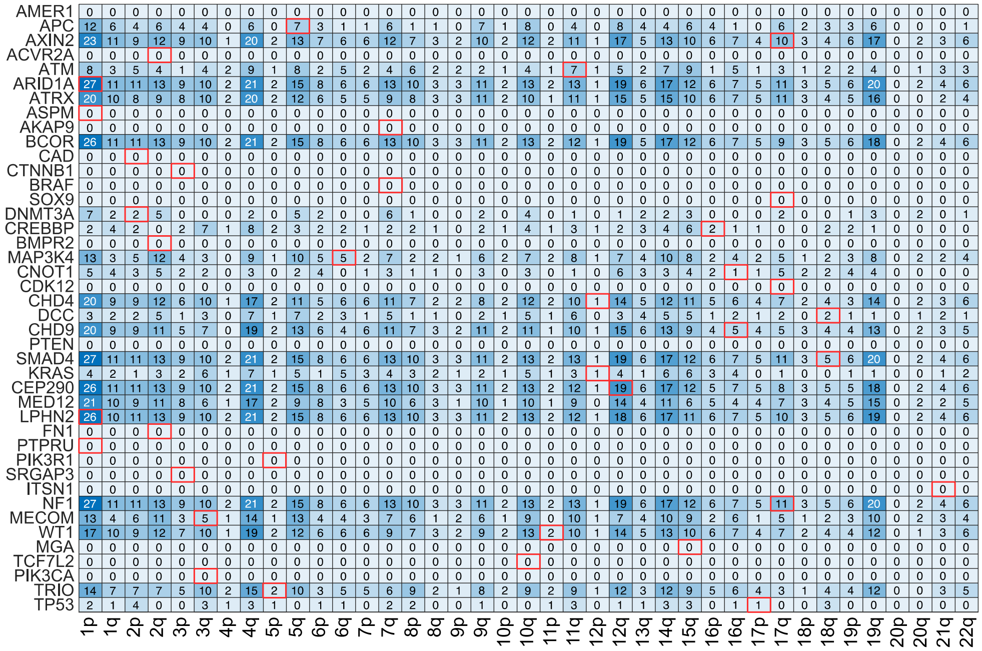


B

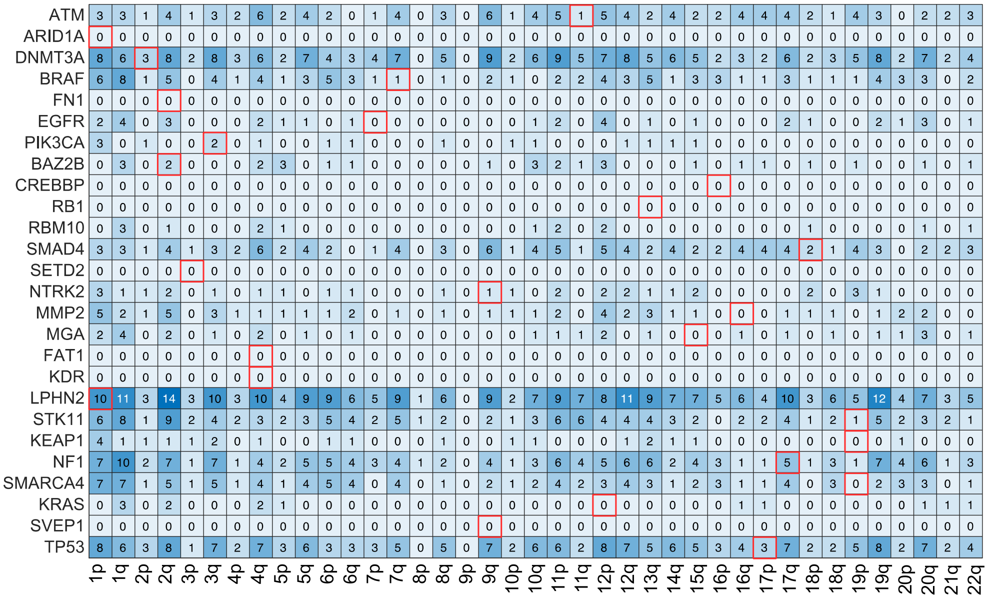

**Figure S9. (A)** 3D scatter plot of the ordered score assigned to the colon training data (X-axis) and the two colon test sets (Y and Z axes). The pairwise rank-correlation coefficients are 0.39 for the training set and test set 1, 0.84 for the training set and test set 2 and 0.41 for the two test sets. All P-values are ~=0. **(B)** Scatter plot of the ordered score assigned to the lung training data (X-axis) and the lung test set (Y axis). The pairwise rank-correlation coefficient is 0.55, with P-value ~=0.

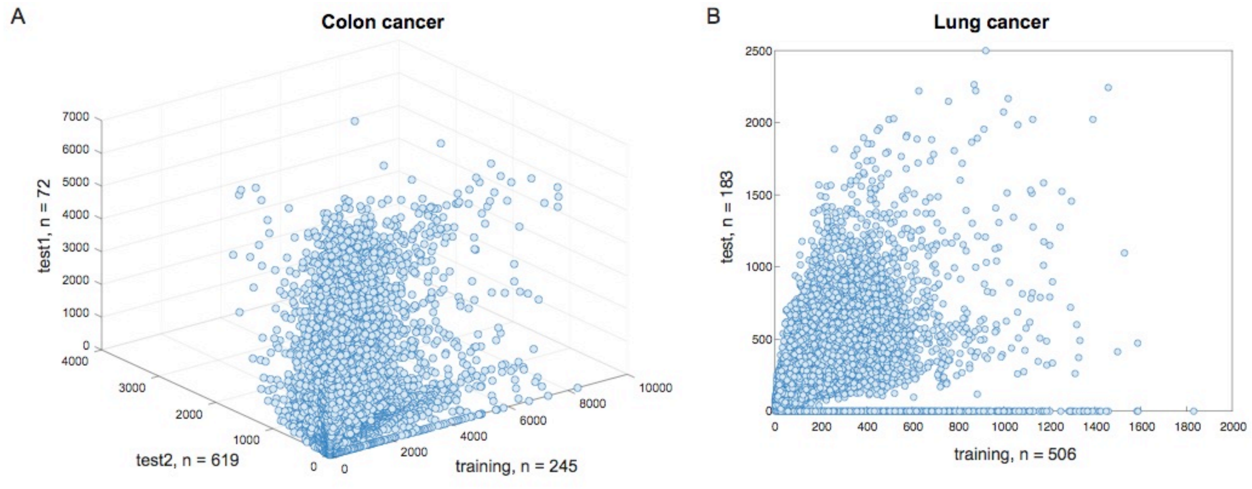

**Figure S10.** Graphical Schema describing utilization of LSTMs to predict interactors of the major drivers.

**(A)** Sequences of occurrences of the major drivers (sequences are of different lengths, from the first ordered driver up to each driver in the sequence) are used to predict the occurrence of other mutations. This is resulting with 42 LSTMs for colon and 26 for lung (the number of major drivers for each tumor type) that are trained to predict the occurrence of other ("passenger") mutations.

(B) Each "Passenger" gene A in which mutations could be predicted with good performance (AUC>0.85) using the test set repeatedly, from multiple locations in the sequence of drivers, is selected as predicted interactor (as it can be robustly predicted from a landscape of major drivers)

(C) To pair A with major drivers (those with which A is predicted to interact), the LSTM scores predicting the occurrence of A from the drivers sequences with high accuracy are utilized. These score are correlated with the occurrence of each major driver. The major drivers that are significantly correlated with the scores predictive of mutations in A are paired with it, as these are likely to be contributing to its prediction.

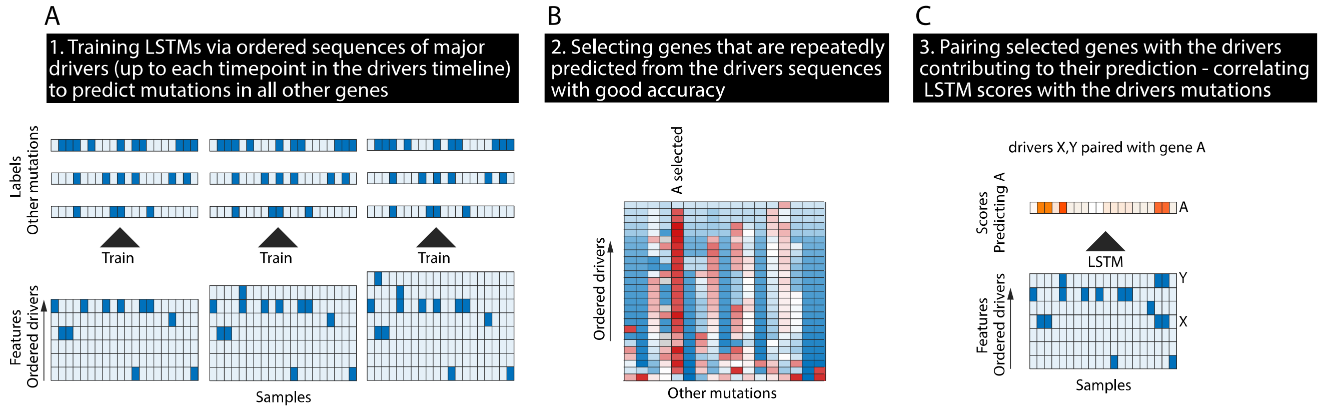

**Table S1.** Performance (AUC) of the prediction of mutations occurrences in the sequence

**Table S2.** Colon cancer reconstructed mutational data

**Table S3.** Lung cancer reconstructed mutational data

**Table S4.** Performance (AUC) predicting the occurrences of driver-interactors from the sequence of drivers

**Table S5.** Pairs of driver-interactor genes

**Table S6.** GO pathways and the enrichment P-values for the joint drivers between colon and lung cancer, and the colon and lung interactors

**Table S7.** Lists of driver genes for colon and lung cancer
